# MASH-in-a-dish reveals fatty acid-driven itaconate crosstalk between Kupffer cells and hepatocytes

**DOI:** 10.64898/2026.10.31.763289

**Authors:** Jung-Chin Chang, Annika Sullock Enzlin, Maya W. Haaker, Ennis Maiguashca Sorensen, Tessa J. van der Leek, Bianca S. Mascarenhas, Thomas Alexander Kluiver, Richard W. Wubbolts, Saskia van Essen, Laura Bongiovanni, Rachel Thomas, Weng Chuan Peng, Coen C. Paulusma, Esther A. Zaal, Bart Westendorp, Alain de Bruin, J. Bernd Helms, Celia R. Berkers

**Affiliations:** Division Cell Biology, Metabolism & Cancer, Department of Biomolecular Health Sciences, Faculty of Veterinary Medicine, Utrecht University, Utrecht, The Netherlands; Institute of Virology, Marburg University, Marburg, Germany; Princess Máxima Center for Pediatric Oncology, Utrecht, The Netherlands; Department of Veterinary Medicine, University of Teramo, Teramo, Italy; Tytgat Institute for Liver and Intestinal Research, Amsterdam Gastroenterology Endocrinology Metabolism (AGEM) Research Institute, Amsterdam UMC, Amsterdam, The Netherlands

**Keywords:** MASH, MASLD, steatosis, inflammation, fibrosis, itaconate, metabolic crosstalk

## Abstract

Metabolic dysfunction-associated steatohepatitis (MASH) is driven by interactions between metabolically stressed hepatocytes and non-parenchymal liver cells, leading to hepatocyte injury, inflammation, and fibrosis. Metabolites can mediate intercellular communication, yet their contribution to MASH pathogenesis remains poorly understood, partly because tissue dissociation disrupts the local metabolic environment. To address this challenge, we developed a novel *in vitro* MASH model, “MASH-in-a-dish”, a Transwell co-culture system of mouse hepatocyte-derived organoids (HepOrgs), Kupffer cells (KCs), and hepatic stellate cells (HSCs) in a serum-free, chemically defined liver co-culture medium. MASH-in-a-dish modelled key features of clinical MASH, including steatosis, hepatocyte ballooning, inflammation and fibrosis, and allowed for rapid cell separation and cell type-specific metabolomics and stable-isotope fluxomics analysis. Using MASH-in-a-dish, we identified 21 MASH-associated interactive metabolites originating from 6 metabolic pathways, including the tricarboxylic acid cycle and the urea cycle. As a proof of concept, we further characterized itaconate-mediated intercellular crosstalk. Under MASH conditions, fatty acids replaced glucose to drive carnitine palmitoyltransferase I (CTP1A)-dependent itaconate production in KCs. Itaconate exacerbated oxidative stress, steatosis, and hepatocyte injury in lipid-stressed HepOrgs. Inhibiting itaconate production by KC-specific *Acod1* knockout in the MASH-in-a-dish system mitigated hepatocyte damage. These findings identify fatty acid-driven itaconate production as a mechanism for lipid stress-induced immunometabolic crosstalk, linking adipose tissue insulin resistance to hepatocyte injury in MASH. In conclusion, our results establish MASH-in-a-dish as a powerful and versatile *in vitro* platform to investigating cell-cell interactions and metabolite crosstalk in MASH pathogenesis, and identify fatty acid-driven itaconate crosstalk between Kupffer cells and hepatocytes as a novel pathogenic mechanism and a potential therapeutic target in MASH.

## 1. Introduction

Affecting nearly 40% of the global population, metabolic dysfunction-associated steatotic liver disease (MASLD) has become the most common liver disease and one of the leading cause of hepatocellular carcinoma (HCC) and liver transplantation^1^. Each year 0.3-2.2% of MASLD patients progress from benign steatosis to metabolic dysfunction-associated steatohepatitis (MASH), which increases the risk for HCC by 27.5 to 110 times^2^. The mainstay of MASLD treatment rely on hypocaloric dietary intervention and life-style modification with 5-10% weight loss as the primary target^3^. While several promising pharmacological agents are in phase 3 clinical trials, Resmetirom (thyroid hormone receptor β-selective agonist) and Semaglutide (glucagon-like peptide-1 receptor agonist) are currently the only approved pharmacotherapies for MASH. Current therapies are still insufficient to curb the MASLD pandemic^1,4^, and new therapeutic targets to mitigate MASLD-related liver failure and HCC are urgently needed.

MASH is characterized by steatosis and ballooning degeneration of hepatocytes, sterile inflammation and fibrosis in a steatotic, metabolically stressed liver^5^. The cellular crosstalk between steatotic hepatocytes, Kupffer cells (KCs), and hepatic stellate cells (HSCs) is central to the initiation and progression of MASH^6^. For instance, crosstalk signaling via proteinaceous ligands, such as hedgehog ligands^7,8^ and interleukin-6 (IL-6)^9^, has been shown to promote the MASH phenotype and carcinogenesis, respectively. However, targeting pro-inflammatory cytokines has not been proven effective clinically and the current MASH therapeutic landscape focuses on correcting the underlying metabolic dysfunctions^10^. In addition to proteinaceous ligands, metabolites are now recognized not only as a source of energy and building block, but also as signaling molecules and novel therapeutic targets^11^. Several metabolites originating from the tricarboxylic acid (TCA) cycle (e.g. itaconate^12–15^, mesaconate^16^, succinate^17^) regulate immune functions in animal models of sepsis and immune-mediated inflammatory diseases. Uridine 5’-diphosphate (UDP)-glucose and UDP-galactose from dying hepatocytes can act as damage-associated molecular patterns (DAMPs) to promote HSC activation and fibrogenesis^18^. Given the role of metabolic dysfunctions in driving sterile inflammation and fibrosis in MASH, metabolite crosstalk in liver likely plays an important role in MASH pathogenesis ^19^. However, how metabolite signaling contributes to MASH initiation and progression remains poorly understood.

Studying metabolite crosstalk in a MASH liver is challenging. Firstly, the identification of cross-talking metabolites *in vivo* or *ex vivo* is difficult because the crosstalk metabolites are present in the interstitial fluid, which is inevitably lost or contaminated with blood metabolites during isolation. Direct extraction of liver tissues yields a mixture of metabolites from liver cells, serum, and blood cells, which is not informative on metabolite crosstalk. Secondly, while separating liver cell types can give insight into cell type-specific metabolic processes, the lengthy procedure needed to separate different liver cell types prior to metabolic profiling drastically distorts metabolism. *In vitro* modelling of MASH can overcome some of these challenges. MASH has been modelled *in vitro* by multi-cellular liver spheroids^20,21^, human pluripotent stem cell-derived organoids^22^, and liver-on-chip^23^. However, these models do not allow rapid separation of the constituent cell types, making them unsuitable for metabolic interaction studies.

To investigate the metabolite crosstalk in MASH liver, we here developed “MASH-in-a-dish”, a Transwell co-culture system of mouse hepatocyte-derived organoids (HepOrgs), KCs, and HSCs. MASH-in-a-dish not only recapitulates the pathological hallmarks of MASH, but is also unique in that it enables rapid separation of the constituent cell types, allowing for, for the first time, cell-type-specific metabolome analyses in a co-culture system. By combining MASH-in-a-dish and liquid chromatography-mass spectrometry (LC-MS)-based metabolomics and stable isotope tracing, we provide evidence that fatty acid-driven hepatic itaconate crosstalk is a promising target for developing novel MASH treatments.

## 2. Results

### 2.1 MASH-in-a-dish models steatotic hepatocyte injury, inflammation, and fibrogenic responses of the MASH liver

Interactions between parenchymal (i.e. hepatocytes) and non-parenchymal liver cells (e.g. KCs and HSCs) underlie the progression of key MASH pathologies, including steatosis, hepatocyte ballooning, inflammation, and fibrosis. To understand lipid-driven metabolite crosstalk between parenchymal and non-parenchymal cells in MASH liver, we developed the MASH-in-a-dish system. MASH-in-a-dish consists of primary mouse hepatocyte-derived organoids (HepOrgs)^24^ in a basement membrane extract (BME) droplet hanging underneath the Transwell membrane, primary mouse Kupffer cells (KCs) in the bottom well, and mouse hepatic stellate cells (HSCs) in the Transwell insert (**Figure 1A**). These cells are cultured in a serum-free, chemically defined liver co-culture medium (L-CCM), which circumvents the interference of undefined metabolites from other cell culture supplements, such as fetal bovine serum. Finally, MASH is induced in this system by a lipid loading mixture formulated for this purpose.

**Figure 1.**
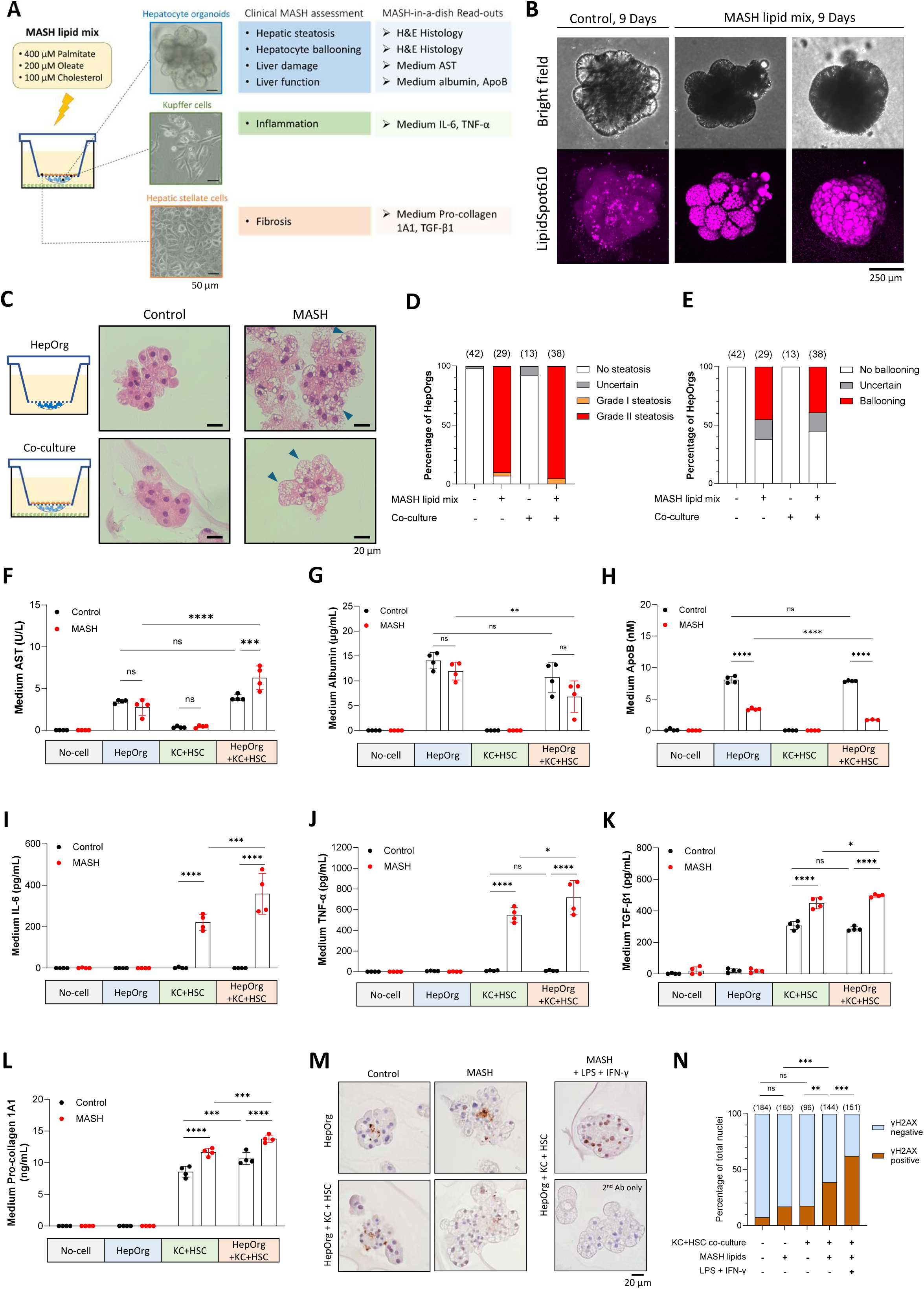
MASH-in-a-dish models steatotic hepatocyte injury, inflammation, and fibrogenic responses of the MASH liver. **(A)** Configuration of the MASH-in-a-dish Transwell co-culture system of primary hepatocyte organoids (HepOrgs), Kupffer cells (KCs), and hepatic stellate cells (HSCs) and the *in vitro* readouts for assessing key clinical presentation of MASH. **(B)** HepOrgs were co-cultured with HSCs in control or MASH medium. Lipid droplets in HepOrgs were visualized with neutral lipid probe LipidSpot610. **(C-E)** Representative pictures and pathological scoring of hematoxylin-eosin staining of HepOrgs cultured alone or with non-parenchymal cells (KC + HSC) in the MASH-in-a-dish system in control medium or MASH medium for 4 days. Total number of scored organoids is indicated in the parenthesis above each condition. Arrow heads indicates ballooning-like cells. **(F-L)** HepOrgs and non-parenchymal cells (KCs + HSCs) were cultured separately or co-cultured in control or MASH medium for 3 days. Aspartate transaminase (AST) activity (F), albumin (G), apolipoprotein B (ApoB, as a readout for VLDL) (H), interleukin 6 (IL-6) (I), tumor necrosis factor α (TNF-α) (J), transforming growth factor β1 (TGF-β1) (L), and pro-collagen 1A1 (L) in the media were determined. Data are presented as mean ± SD of 4 independent MASH-in-a-dish Transwell cultures. **(M)** Representative immunohistochemistry images of γH2AX of HepOrgs cultured alone or with non-parenchymal cells (KC + HSC) in MASH-in-a-dish system in control medium or MASH medium for 4 days. MASH + 10 ng/mL lipopolysaccharide (LPS) + 25 ng/mL murine interferon-γ (IFN-γ) serves as the positive control. **(N)** Pathology scoring of nuclear γH2AX staining in (M); total number of scored nuclei are indicated in the parenthesis above each condition. **Statistical analysis:** (C-I) Two-way ANOVA with Tukey’s multiple comparisons test. (M) χ^2^ test with Bonferroni correction for multiple comparisons. ns, not significant; *, *P* < 0.0332; **, *P* < 0.0021; ***, *P* < 0.0002; ****, *P* < 0.0001.

We first investigated cell viability and liver-specific functions of HepOrgs, KCs, and HSCs in L-CCM before assembling the MASH-in-a-dish co-culture. HepOrg, KCs, and HSCs maintained their viability in L-CCM for at least 3 weeks as evidenced by their stable cell numbers and metabolic activities (**Figure S1A and S1B**). We validated that HepOrgs preserved key lipid metabolic functions in L-CCM, including secretion of conjugated primary bile salts (**Figure S1C**) and apolipoprotein B as a marker for very-low-density lipoproteins (**Figure S1D**). The cultured KCs maintained the KC identity as evidenced by the expression of CLEC4F (KC-specific marker) and F4/80 (macrophage marker) (**Figure S1E and S1F**) and produced pro-inflammatory cytokines IL-6 and TNF-α upon lipopolysaccharide (LPS) stimulation (**Figure S1G and S1H**). Finally, the HSCs retained their characteristic surface stiffness-dependent activation^25,26^, as shown by a significant reduction of pro-collagen 1A1 and TGF-β1 secretion when cultured in a low-stiffness Transwell insert as opposed to a full activation state when cultured on a hard tissue-culture-treated polystyrene surface (**Figure S1I and S1J**).

To induce hepatic steatosis, we designed a MASH lipid mix based on the plasma and liver lipid profiles of metabolic syndrome patients and MASH patients. Patients with metabolic syndrome have increased plasma free fatty acid (FFA), elevated FFA-to-albumin ratio, and total cholesterol, while lipidomes of MASH livers are characterised by increased triglyceride and free cholesterol^27–30^. Based on these characteristics, we formulated a MASH lipid loading mix that consists of 400 µM palmitate, 200 µM oleate, and 100 µM cholesterol. To mimic increased FFA-to-albumin ratio in metabolic syndrome (physiologic range = 0.5 – 2)^30–32^, the fatty acids were prepared as 6:1 bovine serum albumin (BSA) conjugate. Cholesterol was provided as methyl-β-cyclodextrin complex for efficient loading. Upon challenging HepOrgs with MASH lipid mix, we observed increased triglyceride content (**Figure S1K**). In HepOrg mono-culture and HepOrg co-cultured with KCs and/or HSCs, brightfield microscopy showed that the MASH lipid mix induced the development of small (2-200 µm^2^) and large (> 200 µm^2^) lipid droplets^33^ over a course of 9 days (**Figure S1L**). We further confirmed lipid droplets induction by the MASH lipid mix with neutral lipid probes LipidSpot610 (**Figure 1B and Videos S1-S2**) and Bodipy 493/503 (**Figure S1M**) in HepOrgs co-cultured with HSCs. These data showed that hepatic steatosis can be modelled using HepOrgs and our MASH lipid mix.

We next used the fully assembled MASH-in-a-dish to study how the interactions between parenchymal and non-parenchymal cells affect the development of MASH, by comparing HepOrgs that were cultured in absence or presences of KCs and HSCs. In analogy to clinical tests for evaluating the disease activity of MASH and liver functions, we examined clinically-inspired MASH read-outs in MASH-in-a-dish, scoring for steatosis, ballooning hepatocytes, liver function and injury, inflammation and fibrosis (**Figure 1A**). Histological analysis of HepOrgs confirmed that the MASH lipid mix alone was sufficient to induce steatosis and ballooning-like phenotypes in HepOrgs, independent of the presence of KCs and HSCs (**Figure 1C-1E**). Of note, unlike hepatocyte-derived liver organoids, treating intrahepatic cholangiocyte-derived liver organoids with the MASH lipid mix did not result in steatosis (**Figure S1N**).

In contrast to steatosis and ballooning, liver damage, hepatic function, inflammation and fibrosis were synergistically affected by the MASH lipid mix and co-culture with non-parenchymal cells. MASH lipid increased the liver damage marker aspartate transaminase (AST) only in the complete co-culture (**Figure 1F**) and decreased hepatic secretory functions, including secretion of albumin (**Figure 1G**) and very low density lipoprotein (VLDL, **Figure 1H)**. Likewise, the MASH lipid mix induced secretion of proinflammatory cytokine IL-6 and TNF-α, profibrogenic cytokine TGF-β1, and fibrosis marker procollagen 1a1 in the KC-HSC co-culture, which was further enhanced in the presence of HepOrg (**Figure 1I -1L**). Consistent with increased liver injury, inflammatory and fibrogenic responses, co-culture with KCs and HSCs also enhanced DNA damage in hepatocytes, as shown by increased nuclear gH2AX staining (**Figure 1M and 1N**). Together, these results demonstrate that in response to lipid loading, MASH-in-a-dish develops steatosis, hepatocyte injury, inflammation, and fibrosis that mimics clinical MASH.

### 2.2 MASH-in-a-dish reveals metabolic interactions under normal and MASH conditions

The configuration of MASH-in-a-dish allows for rapid separation of individual cell types and thereby rapid extraction of the metabolomes from different cell types for metabolomics and fluxomics analysis. Taking advantage of this property, we investigated the metabolite crosstalk amongst HepOrgs, KCs, and HSCs in the absence or presence of MASH lipid mix. To this end, we performed [U-^13^C6]-glucose tracing for 24 hours using three different co-culture combinations of parenchymal and non-parenchymal cells, followed by LC-MS-based metabolomic profiling of the culture medium and individual cell types within each co-cultures (**Figure 2A**). Principal component analysis (PCA) and unsupervised clustering based on total intracellular metabolite levels revealed that HepOrgs, KCs, and HSC retained their cell-type specific metabolomes in the presence and absence of the MASH lipids (**Figure 2B, Figure S2A**). Of note, metabolomic profiling of MASH-in-a-dish also revealed cell type-specific metabolite synthesis. For instance, bile acids, betaine, and ornithine were predominantly synthesized in HepOrgs while guanidinoacetate and itaconate were specifically synthesized in KCs (**Figure S2A**). PCA for each cell type revealed clear effects of co-culture and MASH treatment on intracellular metabolomes in HepOrgs and KCs, while HSC metabolomes seemed to respond predominantly to MASH lipid treatment (**Figure S2B-S2D**).

**Figure 2.**
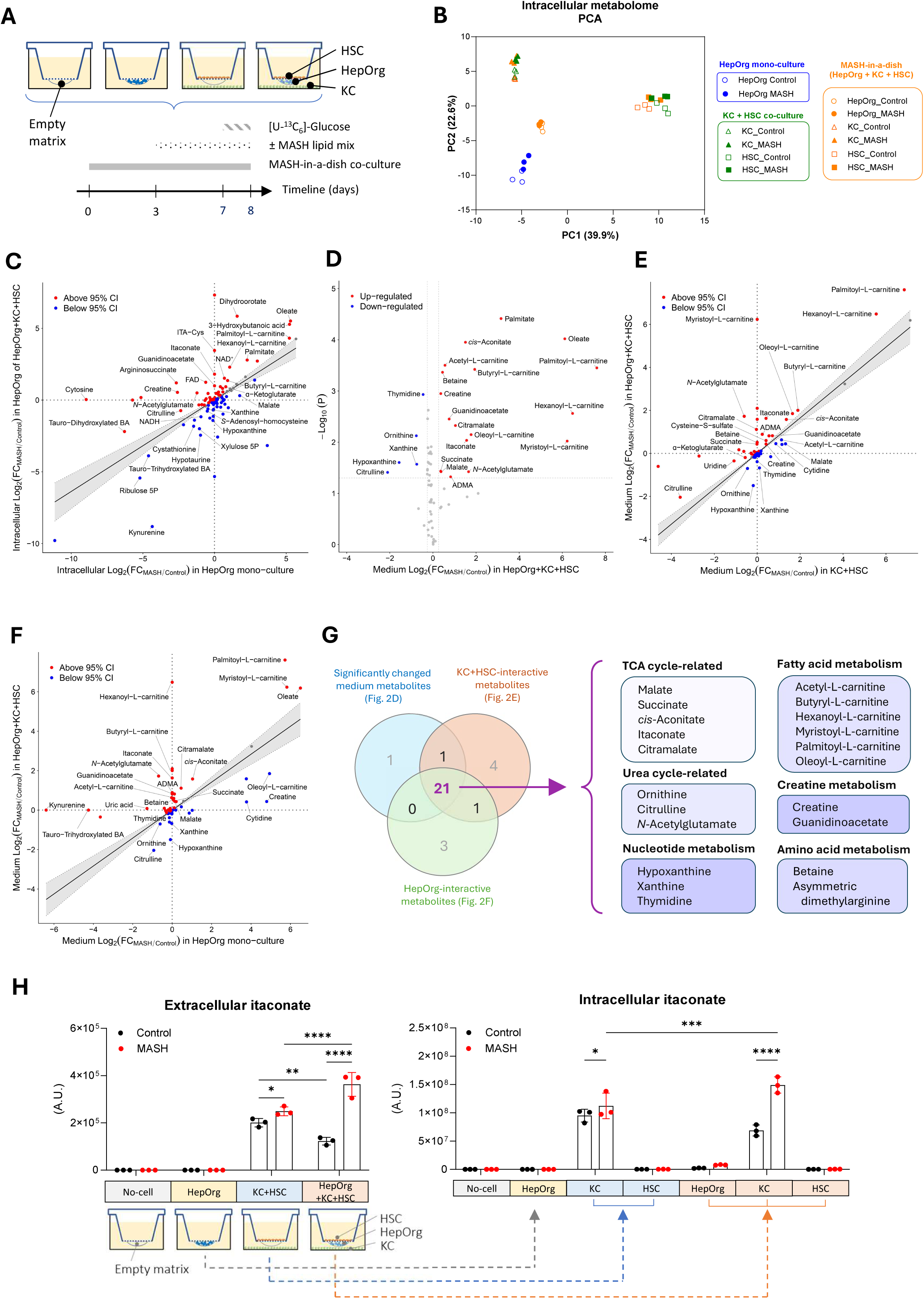
MASH-in-a-dish reveals metabolite crosstalk under normal and MASH conditions. **(A)** Experimental design and timeline for metabolomics and ^13^C6-glucose fluxomics in MASH-in-a-dish. Empty insert control (BME without cells), parenchymal cells alone (HepOrg), non-parenchymal cells alone (KC+HSC), and the complete MASH-in-a-dish (HepOrg+KC+HSC) were cultured in co-culture medium for 3 days and then treated with control medium or MASH medium for 4 days. ^13^C6-glucose was introduced during the last 24 hour of incubation. Both intracellular metabolomes and extracellular (i.e. medium) metabolomes were analyzed. **(B)** Principle component analysis for the intracellular metabolomes for each cell type in different MASH-in-a-dish co-cultures. **(C)** Galaxy plot of intracellular metabolomes of HepOrgs mono-culture versus HepOrgs in MASH-in-a-dish co-cultures (i.e. co-cultured with KC + HSC) treated with MASH medium versus control medium. **(D)** Volcano plots of extracellular metabolomes of MASH-in-a-dish co-cultures treated with MASH medium versus control medium. **(E)** Galaxy plot of extracellular metabolomes of KC + HSC co-culture versus HepOrg + KC + HSC co-culture for identifying HepOrg-interactive metabolites. **(F)** Galaxy plot of extracellular metabolomes of HepOrgs mono-culture versus HepOrg + KC + HSC co-culture for identifying KC+HSC-interactive metabolites. **(G)** Venn diagram for identifying MASH-associated interactive medium metabolites. **(H)** Intracellular and extracellular (i.e. conditioned co-culture medium) levels of itaconate in different MASH-in-a-dish co-cultures; extracted from the metabolomics experiment in (A). n = 3 independent Transwell culture for all conditions. **Statistical analysis:** (H) Two-way ANOVA with Tukey’s multiple comparisons test. *, *P* < 0.0332; **, *P* < 0.0021; ***, *P* < 0.0002; ****, *P* < 0.0001.

As the key driver of MASH phenotypes, we focused on HepOrgs. Volcano plots comparing HepOrgs in the absence and presence of the MASH lipid mixture showed that MASH lipids induce a substantial metabolic rewiring in both single HepOrg and HepOrg+KC+HSC co-cultures (**Figure S2E and S2F**). To identify HepOrg intracellular metabolites whose MASH lipid-induced changes were significantly different in HepOrg+KC+HSC co-culture versus HepOrg monoculture, we projected the MASH-lipid induced Log2 fold change (FC) in these conditions into a “galaxy plot” (**Figure 2C**). In such a galaxy plot, if the presence of non-parenchymal cells has no influence on MASH-induced metabolite changes in HepOrg, then the data point will fall in the 95% confidence interval (CI) of the linear regression line (indicated as a grey area). Conversely, if a MASH-induced metabolite change is influenced by the presence of KC and HSC, the data point will fall outside the 95% CI. This analysis revealed several MASH-induced, cell interaction-responsive metabolites, including itaconate, creatine, citrulline, *N*-acetyl-glutamate, *cis*-aconitate, malate and *N*-carbamoyl-aspartate (**Figure 2C**). Overall, the intracellular metabolome analysis showed that metabolic rewiring takes place in the MASH-in-a-dish system, both as a result of cell-cell interactions within the model and due to the high abundance of lipids.

To identify crosstalk metabolites that can explain the observed metabolome changes in HepOrg in MASH-in-a-dish, we next analyzed the extracellular metabolomes (i.e. medium metabolomes) of the three different co-culture combinations of parenchymal and non-parenchymal cells. We reasoned that a crosstalk metabolite in MASH would fulfil the following criteria: (1) the level of the metabolite in the co-culture medium must either increase or decrease by more than 20% by MASH lipids in HepOrg+KC+HSC co-culture as compared to HepOrg mono-culture, (2) the effect of MASH lipids on the metabolite level in the co-culture medium is dependent on both the presence of HepOrg and the presence of KC+HSC. Volcano plot analysis revealed 23 metabolites that were significantly up-or down-regulated (criterion 1, **Figure 2D**). To identify HepOrg- and KC+HSC-interactive MASH metabolites, we again used galaxy plots, comparing MASH-lipid induced Log2 FC in medium metabolite levels in KC+HSC co-culture versus HepOrg+KC+HSC (criterion 2, **Figure 2E**) as well as in HepOrg mono-culture versus HepOrg+KC+HSC co-culture (criterion 2, **Figure 2F**). This analysis resulted in the identification of 27 KC+HSC-interactive MASH metabolites and 25 HepOrg-interactive MASH metabolites **(Figure 2E and 2F)**. Combining the criteria, we identified 21 MASH-associated interactive metabolites, originating from 6 distinct metabolic pathways: the TCA cycle, fatty acid metabolism, the urea cycle, creatine metabolism, nucleotide metabolism, and amino acid metabolism (**Figure 2G and S2G**).

To investigate which cell type(s) produced the putative cross-talk metabolites, we next looked into the intracellular levels of these 21 metabolites (**Figure S2H**) and observed that in MASH conditions, HepOrgs are mainly responsible for secretion of citramalate and malate, HSCs for medium- and long-chain acyl-carnitines, and KCs for guanidinoacetate, ADMA, itaconate, succinate, cis-aconitate, and short-chain acyl-carnitines. Amongst the 21 candidate metabolites is itaconate, a myeloid lineage-specific immunometabolite that was shown to be highly elevated in human and mouse MASH livers^34,35^. Since liver is the main organ responsible for itaconate degradation^36^, we took advantage of the MASH-in-a-dish model to explore the role of itaconate crosstalk between KCs and HepOrgs in MASH pathogenesis. By combining the intra- and extra-cellular metabolomics data of the MASH-in-a-dish, we found that itaconate is specifically produced by KCs (**Figure 2H**, right panel) and was significantly induced when KCs were co-cultured with HSCs and HepOrgs under MASH conditions (**Figure 2H**, right panel), resulting in increased levels of extracellular itaconate (**Figure 2H**, left panel). In contrast, in the absence of MASH lipids, co-culture of KC+HSC with HepOrgs decreased the itaconate level in the medium, suggesting itaconate is degraded by co-cultured HepOrgs, presumably to citramalate. Indeed, in the presence of MASH lipids, both the intracellular citramalate in HepOrgs as well as extracellular citramalate in the medium were increased in HepOrg+KC+HSC co-culture (**Figure S2I**). Consistent with itaconate being an inhibitor of succinate dehydrogenase, we also observed increased succinate levels in Kupffer cells and in the medium of MASH-in-a-dish (**Figure S2G and S2H**). Together, these observations establish MASH-in-a-dish as a novel and powerful *in vitro* system that enables the dissection of cell-type-specific metabolic contributions to MASH pathogenesis. Using MASH-in-a-dish, we identified a set of MASH-associated crosstalk metabolites, including the immunometabolite itaconate, whose synthesis is increased in KCs while its degradation is enhanced in HepOrgs under MASH conditions.

### 2.3 Fatty acids replace glucose for increased itaconate synthesis in Kupffer cells in MASH

We next asked how the MASH condition drives itaconate production by examining the [U-^13^C6]-glucose labelling patterns of itaconate in primary KCs in the same metabolomics dataset of the completely assembled MASH-in-a-dish presented in Figure 2 (**Figure 3A**). In the absence of MASH lipids, glucose contributed to 46% of all synthesized itaconate with 28% of the itaconate containing one [U-^13^C6]-glucose-derived carbon (M+1) and 18% containing two [U-^13^C6]-glucose-derived carbons (M+2) (**Figure 3B**). M+1 itaconate is consistent with incorporation of M+2 acetyl-CoA into citrate and cis-aconitate and subsequent decarboxylation by cis-aconitate decarboxylase 1 (ACOD1), while M+2 itaconate is likely produced via reverse TCA cycle flow from M+2 succinate in mitochondria where succinate dehydrogenase is suppressed by itaconate. Surprisingly, in the presence of MASH lipids, despite a clear increase in itaconate production, the production of itaconate from glucose was greatly suppressed.

**Figure 3.**
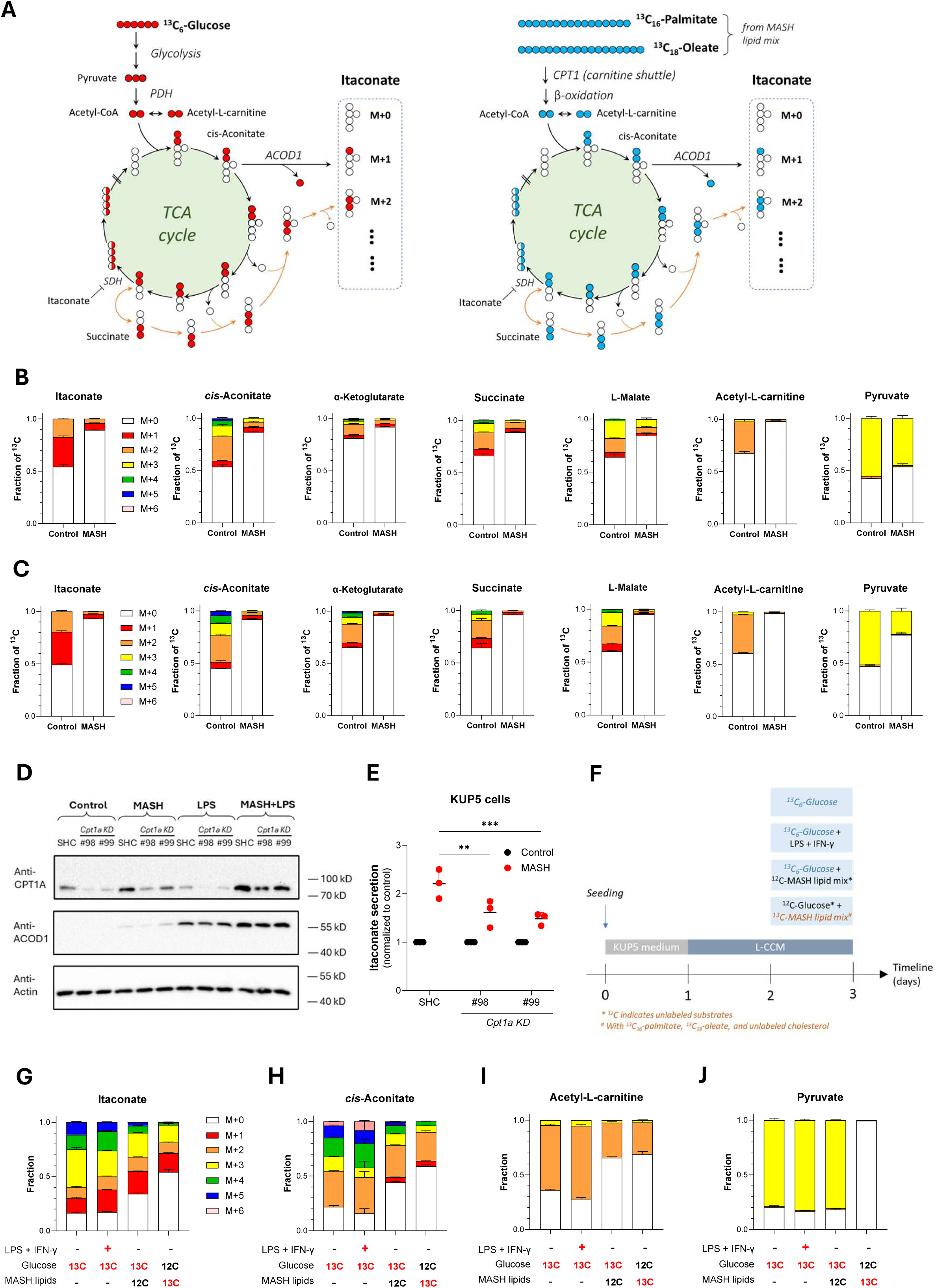
Fatty acids replace glucose for increased itaconate synthesis in Kupffer cells in MASH. **(A)** Schematics for [U-^13^C6]-Glucose tracing (left) and [U-^13^C16]-palmitate/[U-^13^C18]-oleate tracing (right) of itaconate synthesis in Kupffer cells (KCs). Glucose contributes to itaconate synthesis through pyruvate, the glycolysis end-product, while palmitate and oleate contribute to itaconate synthesis through formation of acetyl-CoA and requires carnitine shuttle and mitochondrial fatty acids β-oxidation. **(B-C)** ^13^C-labelled fractions for itaconate, cis-aconitate, α-ketoglutarate, succinate, L-malate, acetyl-L-carnitine, and pyruvate in primary KCs of HepOrg + KC + HSC co-culture (B) and KC + HSC co-culture (C). Data derived from the experiment of Figure 2A. n = 3 independent Transwell cultures. **(D)** Immunoblotting of CPT1A, ACOD1, and total actin (as loading control) in WT, short hairpin control (SHC), and *Cpt1a*KD (#98 and #99) KUP5 cells incubated with control, MASH lipid mix, 100 ng/mL LPS, and MASH + 100 ng/mL LPS for 24 hours. **(E)** LC-MS determination of medium itaconate levels of 3 independently generated polyclonal KUP5 lines of SHC and *Cpt1a*KD (#98 and #99). **(F)** Treatment scheme for [U-^13^C6]-Glucose and [U-^13^C16]-palmitate/[U-^13^C18]-oleate of KUP5 cells in control, LPS + IFN-γ, and MASH conditions. **(G-J)** ^13^C-labelled fractions for itaconate, *cis*-aconitate, acetyl-L-carnitine, and pyruvate in WT KUP5 cells treated as in (F). n = 4 independent Transwell cultures. **Statistical analysis: (E)** Two-way ANOVA with Dunnett’s multiple comparisons test. **, *P* < 0.0021; ***, *P* < 0.0002. **Abbreviations:** PDH, pyruvate dehydrogenase; ACOD1, cis-aconitate decarboxylase; SDH, succinate dehydrogenase ; CPT1A, carnitine palmitoyltransferase 1A.

To further investigate itaconate production in the presence of MASH lipids, we next examined [U-^13^C6]-glucose labelling patterns in metabolites of the tricarboxylic acid (TCA) cycle, from which itaconate is derived. The [U-^13^C6]-glucose labelling fractions of many TCA cycle metabolites (cis-aconitate, α-ketoglutarate, succinate, L-malate) and acetyl-carnitine in primary KCs in HepOrg+KC+HSC co-culture (**Figure 3B**), which can be used as proxy for acetyl-CoA, were significantly decreased by MASH lipids while total levels of most of these metabolites were increased (**Figure S3A**). Of note, the [U-^13^C6]-glucose labelling fractions labelling of pyruvate was less affected by MASH treatment, suggesting that an alternative carbon source fuels itaconate production via acetyl-CoA (**Figure 3B)**. Although the MASH lipid mix did not significantly increase the total intracellular levels of itaconate, most TCA cycle metabolite, pyruvate and acetyl-L-carnitine in primary KCs in KC+HSC co-culture (i.e. in the absence of HepOrgs) (**Figure S3B**), we still observed a significant reduction in [U-^13^C6]-glucose labelling of itaconate, TCA cycle metabolites, acetyl-L-carnitine, and pyruvate (**Figure 3C**). These observations showed that KCs used a non-glucose source for itaconate synthesis in MASH conditions. We hypothesized that under MASH conditions, KCs import fatty acids as a major non-glucose carbon source to drive synthesis of itaconate. Because fatty acids must be imported into the mitochondrial matrix via the carnitine shuttle before they can contribute to itaconate synthesis, we tested whether knockdown of carnitine palmitoyltransferase 1A (*Cpt1a*) of the carnitine shuttle can reduce itaconate synthesis in KCs under the MASH condition. Because it is challenging to perform genetic modification and selection in primary KCs, we knocked down carnitine *Cpt1a* in KUP5 cells, a c-myc-immortalized murine Kupffer cell line^37^, by lentivirus-mediated short hairpin expression (**Figure 3D**). We observed that *Cpt1a* knockdown with two different short hairpin RNA constructs significantly reduced itaconate production induced by the MASH lipid mix (**Figure 3E**), suggesting that extracellular long-chain fatty acids are potential carbon sources for itaconate production under MASH conditions.

To test if long-chain fatty acids in the MASH lipid mix could replace glucose as the carbon source for itaconate synthesis in the MASH condition, we performed stable isotope tracing with [U-^13^C6]-glucose versus [U-^13^C16]-palmitate/[U-^13^C18]-oleate in immortalized murine Kupffer cells KUP5^37^ and used LPS + IFN-γ as a positive control for itaconate induction (**Figure 3F**). At the total intracellular metabolite levels, both the MASH lipid mix and LPS + IFN-γ induced itaconate levels in KUP5 cells as expected (**Figure S3C**), but only the MASH lipid mix increased pyruvate levels (**Figure S3D**). In contrast, LPS + IFN-γ treatment significantly increased the level of acetyl-L-carnitine but decreased the level of cis-Aconitate (**Figure S3E and S3F**). We next examined the [U-^13^C6]-glucose labelling of itaconate in KUP5 cells. At the resting state, the [U-^13^C6]-glucose labelling fractions of itaconate, cis-Aconitate, acetyl-L-carnitine, and pyruvate are overall higher in c-myc-immortalized KUP5 cells than in the primary KCs, consistent with c-myc-mediated metabolic rewiring^38^ (**Figure 3G-3J**). In both resting and LPS+IFN-γ-activated KUP5 cells, [U-^13^C6]-glucose labelled about 87% of itaconate. In contrast, in MASH-activated KUP5 cells, the [U-^13^C6]-glucose labelling fraction of itaconate was lowered to 66% (**Figure 3G**), reproducing the finding in primary KCs (**Figure 3B and 3C**). Similarly, MASH lipid mix, but not LPS+IFN-γ reduced the [U-^13^C6]-glucose labelling fractions in cis-aconitate and acetyl-L-carnitine (**Figure 3H and 3I**). Together with the unchanged [U-^13^C6]-glucose labelling in pyruvate (**Figure 3J**), this suggests that MASH-activated KCs produces itaconate from an alternative source of acetyl-CoA, likely fatty acids. Indeed, [U-^13^C16]-palmitate/[U-^13^C18]-oleate tracing confirmed that long-chain fatty acid carbons directly contributed to the synthesis of itaconate, cis-aconitate, and acetyl-L-carnitine under MASH conditions (**Figure 3G-3I**). These data showed that, under MASH conditions, fatty acid-derived acetyl-CoA replaces glucose-derived acetyl-CoA as a primary carbon source for itaconate synthesis by KCs.

### 2.4 Itaconate crosstalk from Kupffer cells promotes steatotic hepatocyte injury

We finally investigated whether itaconate, as a MASH-associated crosstalk metabolite, affects MASH phenotypes in hepatocytes. Itaconate is a nucleophilic metabolite and is known to affect thiol redox homeostasis by alkylating the sulfhydryl groups of glutathione, free cysteine, and cysteine residues^14,15^ and by promoting mitochondrial reactive oxygen species (ROS) production in macrophages^39,40^. Because hepatocytes are the main liver cell type that takes up itaconate for degradation, we hypothesized that chronic exposure to excessive itaconate increases oxidative stress by disrupting thiol redox homeostasis or by increasing ROS production, leading to increased steatotic damage and decreased hepatocyte functions. Analysis of the metabolomics data presented in Figure 2 confirmed increased formation of itaconylcysteine (i.e. the adduct of itaconate and cysteine) in HepOrgs in MASH-in-a-dish co-culture, suggesting that KC-derived itaconate affects the thiol redox homeostasis of HepOrgs (**Figure S4A**). Next, we treated HepOrg 2D mono-culture on collagen-coated surface with itaconate and found that itaconate treatment promoted ROS production, increased AST release (hepatocyte damage marker), and decreased albumin production (liver function marker) (**Figure 4A-C**). Because the intracellular concentration of itaconate in activated macrophages can reach 5-8 mM^14,41^ and itaconate is actively exported by ATP-binding cassette transporter G2 (ABCG2)^42^, we treated HepOrg with increasing concentrations of itaconate up to 8 mM and found that chronic exposure to itaconate dose-dependently promoted lipid accumulation under MASH conditions (**Figure 4D and S4B**). Together, these data suggest that itaconate contributes to MASH phenotypes in hepatocytes by increasing oxidative stress and lipid accumulation.

**Figure 4.**
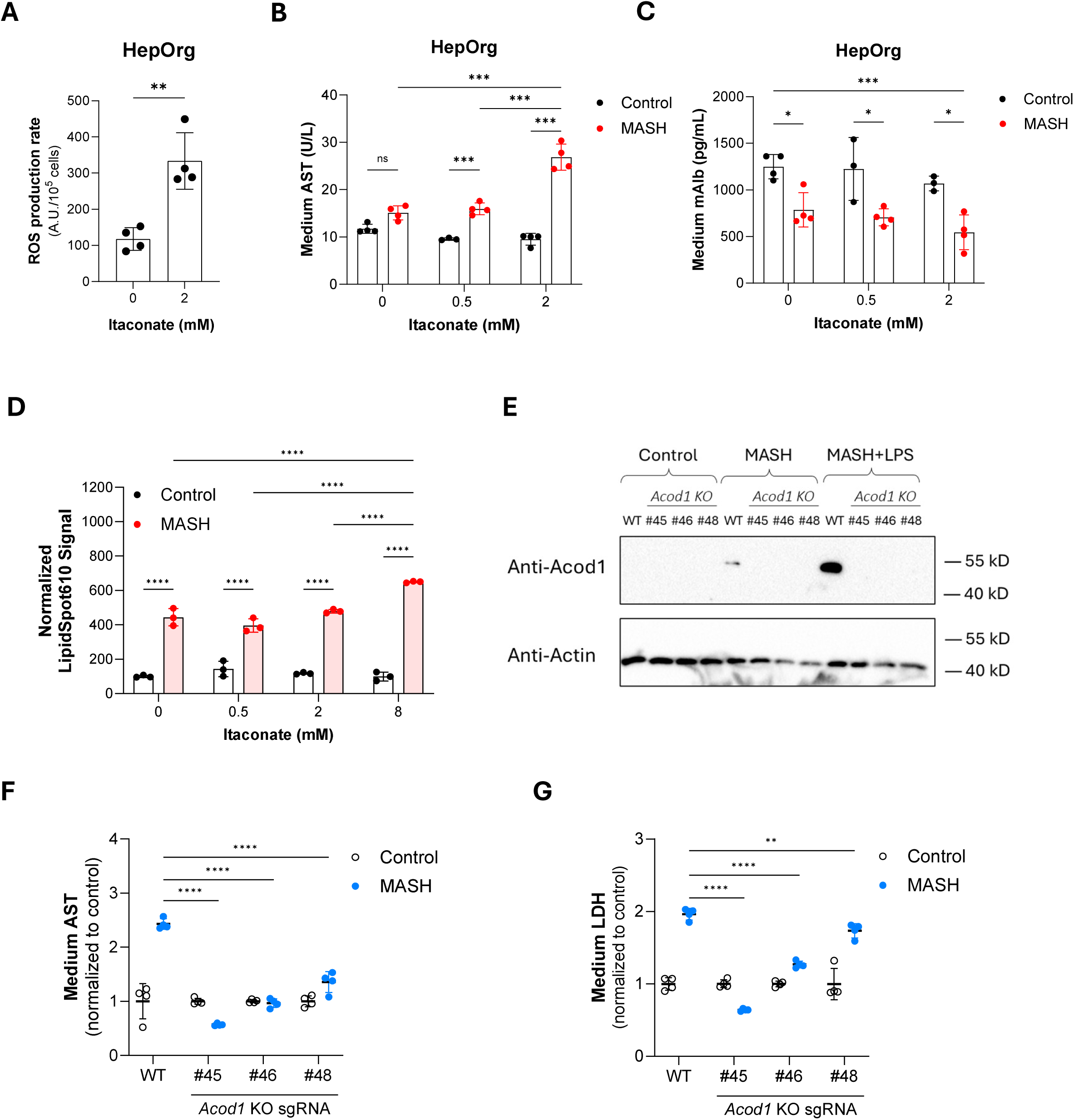
Itaconate crosstalk from Kupffer cells promotes hepatocyte injury. **(A)** HepOrgs were seeded on a rat tail type I collagen-coated surface and treated with 0 or 2 mM itaconate for 4 days. Production rate of reactive oxygen species (ROS) was measured by oxidation rate of dihydroxyethidium in a live cell microplate assay and normalized to cell numbers (as determined by dsDNA content). Data are presented as mean ± SD. Representative data of 3 independent experiments; the incubation condition for itaconate-induced injury varies among organoid lines, ranging from 2 to 8 mM itaconate for 3 to 6 days. **(B-C)** HepOrgs seeded on rat tail type I collagen-coated surface were treated with 0, 0.5, or 2 mM itaconate in control medium or MASH medium for 3 days. Medium AST activity (B) and albumin level (C) were measured. Shown here representative data of 3 independent experiments; n = 4 for (B) and n = 3-4 for (C). The incubation condition for itaconate-induced injury varies among HepOrg lines, ranging from 2 to 8 mM itaconate for 3 to 6 days. **(D)** Effect of itaconate and MASH lipid mix on neutral lipid accumulation in HepOrgs seeded on rat tail type I collagen-coated surface. Neutral lipids were stained with LipidSpot610 and quantified by well scan in a fluorometer and normalized to dsDNA staining with Hoechst33342 from the same well. The data are presented as mean ± SD (n = 3) and is representative of 3 independent experiments. **(E)** Immunoblotting of ACOD1 and total actin (as loading control) for the validation of 3 *Acod1* knockout KUP5 clones generated with sgRNA #45, #46, and #48, respectively. The cells were incubated in control, MASH lipid mix, or MASH lipid mix + 100 ng/mL LPS media for 24 hours. **(F-G)** WT KUP5 and 3 validated *Acod1*KO KUP5 clones were incorporated in the MASH-in-a-dish co-culture with WT HepOrgs and WT HSCs. Medium AST (F) and LDH (G) activities were measured on Day 6 after the MASH lipid challenge. Data are presented as mean ± SD of 4 independent transwell co-culture for KUP5 WT and *Acod1KO* clones. **Statistical analysis:** (A) Two-tailed Student’s *t*-test (B-D) Two-way ANOVA with Tukey’s multiple comparisons test. (F-G) Two-way ANOVA with Dunnett’s multiple comparisons test. ns, not significant; *, *P* < 0.0332; **, *P* < 0.0021; ***, *P* < 0.0002; ****, *P* < 0.0001.

Based on the effect of itaconate on HepOrgs and the observation that KCs are the sole source of itaconate in MASH-in-a-dish (**Figure 2H**), we reasoned that interrupting itaconate crosstalk in MASH-in-a-dish could protect against lipid-induced liver injury. To test this, we knocked out *Acod1* in KUP5 cells by lentivirus-mediated CRISPR-Cas9 delivery with 3 different sgRNAs, performed limiting dilution cloning, and screened for knockout clones by measuring MASH lipid-induced itaconate production by LC-MS (**Figure S4C**). We subsequently validated one knockout clone for each sgRNA by Western blot. In WT KUP5 cells, ACOD1 protein was clearly induced by MASH lipid mix and significantly augmented by combined treatment of MASH lipid mix and LPS. In contrast, neither MASH lipid mix nor MASH lipid mix plus LPS induced detectable ACOD1 expression in three *Acod1*^KO^-KUP5 clones (**Figure 4E**). We then assembled MASH-in-a-dish co-culture with WT KUP5 cells and three independent, validated *Acod1*KO-KUP5 clones and tested whether *Acod1* knockout could rescue MASH phenotypes. While MASH lipid mix challenge induced a 2.4-fold increase in AST release in MASH-in-a-dish with WT KUP5, MASH-in-a-dish with *Acod1*KO-KUP5 clones showed protection against MASH lipid challenge (**Figure 4F)**. Similarly, KC-specific *Acod1* knockout also mitigated the increase of medium LDH (cell death marker) in MASH-in-a-dish (**Figure 4G**). These findings identify itaconate as a MASH-induced crosstalk metabolite that mediates interactions between KCs and HepOrgs and promotes hepatic injury in MASH.

## 3. Discussion

Our present study establishes MASH-in-a-dish as a unique *in vitro* liver model that recapitulates key clinical MASH phenotypes and can be rapidly disassembled for metabolite crosstalk studies. Using a serum-free, chemically defined L-CCM medium and a rationally designed MASH lipid mix, key MASH phenotypes, including steatosis, hepatocyte ballooning and injury, inflammation, and fibrosis, can be robustly induced in this model in 4 days. In contrast, it takes 20-30 weeks for preclinical dietary mouse MASH models to develop MASH^43,44^ and years to decades to develop MASH in humans^2,45^. The significantly faster MASH establishment in MASH-in-a-dish is therefore an advantage that allows mechanistic studies involving cell-cell crosstalk and drug screening. The stability of individual cell types in L-CCM further allows investigation of *in vitro* MASH progression up to three weeks.

By applying metabolomics to the MASH-in-a-dish system, we identified 21 MASH-associated crosstalk metabolites, including the immune-modulatory metabolite itaconate^12,14,15,46^. Several lines of evidence from the present study support itaconate as a crosstalk metabolite in MASH-in-a-dish. First, the MASH lipid mix induced itaconate production in KCs in MASH-in-a-dish and elevated medium levels of itaconate. Increased intracellular levels of itaconate and citramalate in co-cultured HepOrgs further demonstrate that KC-derived itaconate is taken up and metabolized by HepOrgs. Second, treating HepOrgs with itaconate resulted in cell injury, increased ROS production, and increased neutral lipid accumulation. Third, KC-specific knockout of *Acod1* strongly reduced hepatocyte injury in MASH-in-a-dish.

The independent validation of increased levels of itaconate in the liver in dietary-induced murine MASH models^44^ and in livers of MASH patients^34^ strengthens the translatability of MASH-in-a-dish and our metabolomics-based crosstalk study. Consistently, the expression of *Acod1*/*ACOD1* in liver is also upregulated in mice fed with high-fat diet^35,47^ and in MASH patients as compared to non-MASH livers^34,35^. Published *in vitro* and *in vivo* studies further corroborate that itaconate is a tightly regulated crosstalk metabolite in the liver. First, liver mitochondria are known to efficiently oxidize itaconate into CO2^48,49^. Consistent with this, in patients with MASH, itaconate levels are increased specifically in the liver, while remaining unchanged in plasma^34^. Second, *in vivo* itaconate tracing showed that the itaconate degradation pathway is most active in the mitochondria of the liver and kidneys^36^. Third, the protein level of solute carrier SLC13A3, a plasma membrane transporter for itaconate, was elevated in the liver of a murine MASH model induced by a high-fat, high-cholesterol, high-fructose diet. Hepatocyte-specific knockdown of *Slc13a3* in this model significantly reduced liver weight, liver cholesterol level, and liver triglyceride level^50,51^. Fourth, knockout of *Sugct*, the first mitochondrial enzyme involved in itaconate degradation in the liver and kidneys, resulted in steatosis of both organs^52^.

In addition to identifying itaconate as a MASH-associated crosstalk metabolite, we uncovered a novel synthesis route of itaconate in KCs under the MASH conditions using [U-^13^C6]-glucose and [U-^13^C16]-palmitate/[U-^13^C18]-oleate tracing. Previous studies characterised the synthesis routes of itaconate in toll-like receptor 4 (TLR4) agonist-activated murine bone marrow-derived macrophages (mBMDM)^53,54^ and macrophage-like cell lines (e.g. RAW246.7^41^) using various stable isotope tracing strategies. In TLR4-agonist-activated macrophages, tracing with [U-^13^C6]-glucose revealed itaconate synthesis via forward (oxidative) TCA cycle activity, resulting in M+1 and M+3 itaconate. Additionally, [U-^13^C5]-glutamine tracing in LPS-activated RAW264.7 cells, mBMDM, and human monocyte-derived macrophages showed that 35-40% of glutamine-derived itaconate is synthesized via reversed (reductive) TCA cycle activity^55^, likely because of the competitive inhibition of succinate dehydrogenase by itaconate^46^. Here we extend these studies to primary and immortalized KCs, showing that in a non-MASH resting state or under LPS-activated conditions, itaconate is synthesized from [U-^13^C6]-glucose via both the forward and reverse TCA cycle. Additionally, we found that MASH-activated primary KCs synthesize itaconate predominantly from a non-glucose source and showed that MASH-activated immortalized KCs (KUP5 cells) upregulate CPT1A and synthesize significant amounts of itaconate using palmitate/oleate-derived acetyl-CoA in a CPT1A-dependent manner. Given that itaconate can stabilize CPT1A to enhance fatty acid oxidation^56^, our data thus suggest a feedforward mechanism of itaconate synthesis by KCs under MASH conditions, where fatty acids upregulate ACOD1 and CPT1A to drive itaconate synthesis and the increased level of itaconate in turn stabilizes CPT1A to sustain fatty acid import into mitochondria for itaconate synthesis. These findings provide a mechanistic link between increased fatty acids release from insulin-resistant adipose tissue and elevated itaconate and CPT1A in the MASH liver. The pathogenic importance of CPT1A in MASH pathogenesis is further supported by increased CPT1A expression in activated HSCs in MASH livers; inhibition of CPT1A in this context suppresses the fibrogenic response of activated HSCs^57^.

Our findings also indicate that chronic increase of hepatic itaconate levels may exacerbate MASH. We found that itaconate promotes ROS production, lipid accumulation, hepatocyte injury in HepOrgs and that KC-specific knockout of *Acod1* in MASH-in-a-dish co-culture dampens lipid-induced hepatocyte injury. This is in line with previous studies showing that itaconate promotes macrophage ROS production by stimulating NADPH oxidase activity^58^ and fatty acid-dependent mitochondrial ROS production^39^, and suppressing peroxiredoxin 5^40^. The role of itaconate in MASLD development has previously been investigated in diet-induced MASLD mouse models in combination with systemic manipulation of itaconate signaling, either by Acod1 knockout or by administration of itaconate. In a study by Eberhart et al., germline whole body knockout of *Acod1 (Acod1^-/-^)* offered protection against 16-week high-fat diet (HFD)-induced obesity, insulin resistance, and hepatic steatosis in male C57BL/6J mice while itaconate administration in the drinking water aggravated weight gain under HFD^35^. In contrast, Weiss et al. showed that western diet (WD) with 10% sucrose water increased liver triglyceride content in *Acod1^-/-^* C57BL/6J male mice, but not in the WT mice. The discrepancies in the findings across these mouse studies may be explained by variations in the effectiveness of MASH induction. In addition, whole-body knockout of *Acod1* also obliterates itaconate signaling in other tissues. For instance, the lack of antibacterial effects of itaconate^59^ can potentially promote intestinal bacterial translocation and thereby worsen MASH phenotypes^59,60^. In the present study, our MASH lipid mix is similar to the high-fat diet used in the study of Eberhart et al^35^ (i.e. no fructose) and reproduces the hepatoprotective effect of *Acod1* knockout on MASH phenotypes. Moreover, the protective effect of KC-specific *Acod1* knockout in the MASH-in-a-dish system implies that the locally increased itaconate levels in the livers of MASH patients may contribute to hepatocyte damage in MASH.

In addition to being a crosstalk metabolite in the MASH liver, itaconate has also been shown to create an immune-suppressive tumor microenvironment and to promote tumor progression^61–63^. Recently, itaconate uptake via SLC13A3 into lung cancer cells has been shown to confer resistance to immunotherapy by alkylating programmed cell death ligand 1 (PD-L1)^51^. This suggests that the elevated itaconate levels found in MASH liver can contribute to immunosuppression and may explain the accelerated carcinogenesis rate and resistance to immunotherapy in MASH-related hepatocellular carcinoma (HCC) compared to viral hepatitis-related HCC. A recent study by Natarajan et al. further showed that itaconate can also act as an oncometabolite to promote ZFTA-RELA ependymomas^64^. Because MASH-in-a-dish also models increased DNA damage in MASH resulting from cell-cell interactions, MASH-in-a-dish can potentially be employed to investigate the role of cell-cell crosstalk and metabolite signalling in the initiating phase of MASH-associated HCC.

In conclusion, the presented MASH-in-a-dish system models key clinical MASH phenotypes, including steatosis, hepatocyte ballooning, inflammation, and fibrosis. The contact-free co-culture configuration of the MASH-in-a-dish system allows for rapid cell separation and enables metabolic crosstalk studies. Combining MASH-in-a-dish with metabolomics revealed several MASH-associated interactive crosstalk metabolites, including itaconate. Under MASH conditions, itaconate is synthesized in Kupffer cells in a CPT1A-dependent manner, driven by the increased fatty acid load. Through cell-cell crosstalk, itaconate secreted by Kupffer cells induces oxidative stress, steatosis, and cell injury in hepatocytes. Our study provides a proof of concept for the role of metabolite-mediated crosstalk signaling in MASH pathogenesis and identified itaconate crosstalk between Kupffer cells and hepatocytes as a potential therapeutic target for MASH.

### Limitations

There are a few limitations to the presented MASH-in-a-dish model. First, although we successfully model the key MASH phenotype using the MASH-in-a-dish model, cell-cell contacts are lacking in MASH-in-a-dish. Therefore, certain contact dependent signalling (e.g. notch signalling) or close-range cell-cell paracrine signalling are not modelled but may still play a role in MASH pathogenesis. Second, in the MASH-in-a-dish system, the ratio of “interstitial volume” (i.e. medium volume, 800 µL) to the combined intracellular volume of all cellular components (< 10 µL) is > 80, while the interstitium-to-intracellular ratio is about 0.25 in the liver of healthy volunteers^65^. This means that the impact of metabolite signalling is significantly underestimated due to metabolite dilution. Third, while we incorporated Kupffer cells to model liver inflammation, the immune landscape can be further improved by incorporating other immune cells, such as T cells and dendritic cells. Fourth, while we deactivated HSCs by culturing them on the low-stiffness Transwell insert, HSC activation in MASH may be better modelled by replacing cultured HSCs with freshly isolated quiescent HSC, whose quiescent state can be maintained in the BME matrix^25^. Lastly, gut microbiome metabolites, for instance ethanol^66^ or fructose-derived acetate^67^, play a key role in shaping the metabolic microenvironment in liver but are not incorporated in the current MASH lipid mix.

## 4. Materials and Methods

### 4.1 Primary hepatocyte isolation and culture of hepatocyte organoids (HepOrgs)

The animal experiments were approved by the Netherlands Central Authority for Scientific Procedures on Animals (CCD No. AVD11800202010784) and followed the Amsterdam UMC Institutional Welfare Committee guidelines. Primary mouse hepatocytes (PMHs) were isolated from 6 to 10 weeks old C57BL/6 male mice by 2-step collagenase liver perfusion via the portal vein as previously described^68^. To improve the quality of isolated hepatocytes and non-parenchymal cells, the perfusate was supplemented with 1 mM L-lactate, 0.1 mM pyruvate, and 0.5X MEM essential amino acids solution (Gibco 11130051). To establish hepatocyte-derived organoids, the isolated PMHs were resuspended at 9 × 10^5^ cells / mL in basement membrane extract (BME, R&D, 3533-010-02), plated as 20 µL droplets, and cultured in expansion medium and sub-cultured at 1:2 to 1:3 every 2 weeks as described by Peng et al^24,69^. To enhance outgrowth from single cells and reduce cholangiocyte transformation, the expansion medium was supplemented with 1 ng/mL human IL-6, 0.1 µg/mL glucagon, 4 µM palmitate (as 6:1 fatty acid-BSA conjugate), 2 µM oleate (as 6:1 fatty acid-BSA conjugate), 2 mM taurine, 5 µM hypoxanthine, 0.5 µM thymidine, 50 µM choline, 0.25 mM myo-inositol, 0.5 mM sodium pyruvate, 2 mM sodium lactate, 10 mM HEPES-NaOH (pH 7.4), and 1X MEM amino acid solution. In our hands, preparing A83-01 in anhydrous DMSO and supplementing 1 µM A83-01 freshly or re-supplementing 1 µM A83-01 weekly reduce cholangiocyte trans-differentiation. HepOrgs were matured in L-CCM (see 4.7 for details) for at least 3 days before experiments.

### 4.2 Culture of murine intrahepatic cholangiocyte organoids (ICOs)

Murine intrahepatic cholangiocyte organoids were a kind gift from dr. Bart Spee and were maintained in expansion medium in BME as previously described^70^. For steatosis induction, ICOs were treated with control or the MASH lipid mix in L-CCM.

### 4.3 Primary murine hepatic stellate cells (HSCs) isolation and establishment of HSC cell line

The animal experiments were approved by the Netherlands Central Authority for Scientific Procedures on Animals (CCD No. AVD1080020174484 and No. AVD11800202010784) and followed the Institutional Welfare Committee guidelines at Utrecht University and Amsterdam University Medical Centers. Quiescent hepatic stellate cells (qHSCs) were isolated from adult male C57BL/6J wild-type mice as described^71^. Successful isolation of qHSC were confirmed by the presence of large UV-fluorescent lipid droplets. Freshly isolated qHSCs were cultured in DMEM supplemented with 10% FBS, 2 mM glutamine, and 100 U/L penicillin and 100 µg/mL streptomycin on a TC-treated PS surface for spontaneous activation over a course of 2 weeks. The HSCs were passaged to establish an HSC cell line. For MASH-in-a-dish, the HSC cell line was adapted to William’s medium E (the base medium for MASH-in-a-dish co-culture) and passaged twice weekly.

### 4.4 Primary murine Kupffer cells (KCs) isolation and culture

Primary murine KCs were isolated from the non-parenchymal cells (NPCs) from the leftover cells following hepatocyte isolation^68^ or hepatic stellate cell isolation^71^. To obtain NPCs, the liver cells centrifuged at 50 *× g* for 5 minutes at 4°C to deplete hepatocytes. The supernatant was harvested and centrifuged at 500 *× g* for 5 minutes at 4°C to pellet NPCs. The NPCs from 1 mouse liver were resuspended in seeding medium (William’s medium E supplemented with 10% [v/v] FBS, 100 U/L penicillin and 100 µg/mL streptomycin, 100 µg/mL Normocin, and 2 mM Glutamax) and seeded on a 145 mm non-cell-culture (TC)-treated culture dish. The NPCs were allowed to attach for 4 hours to overnight. Kupffer cells were selected by their strong adhesion to non-cell-culture treated culture dish by washing 5 times with 25 mL cold PBS (pH 7.4) supplemented with 1 mM EDTA. To minimize potential contamination of monocytes, the cells were starved 1 week of M-CSF by culturing in the seeding medium. Afterwards, the culture medium was switched to a in-house Kupffer cell medium (KCM). The KCM is a Williams’ medium E supplemented with 10% (v/v) FBS, 2 mM Glutamax, 50 µg/mL Normocin, 0.1% (w/v) Poloxamer 188, 5 mM Taurine, 100 µM choline chloride, 20 µM hypoxanthine, 1 µM thymidine, 10 ng/mL murine M-CSF. Primary murine KCs were sub-cultured at 70% to 80% confluency by TrypLE supplemented with 0.1% (w/v) Poloxamer 188 and subsequently scraping in PBS supplemented with 1 mM EDTA and 0.1% (w/v) Poloxamer 188. Primary murine KCs were sub-cultured at 1:2 to 1:3 split ratio when reaching 70% to 80% confluency. With this method, we can reliably culture primary murine KCs in vitro for about 2 months.

### 4.5 Cell culture of murine Kupffer cell line KUP5

The c-myc-immortalized murine Kupffer cell line KUP5 was obtained from RIKEN cell bank (#RCB4627)^37^. The cells were cultured in Williams’ medium E supplemented with 10% (v/v) FBS, 100 U/L penicillin and 100 µg/mL streptomycin, 2 mM Glutamax, 10 µg/mL bovine insulin, and 250 µM 1-thioglycerol. The KUP5 cells were cultured on non-TC sub-cultured dishes and sub-cultured twice a week using TrypLE at a split ratio of 1:8 to 1:16.

### 4.6 Preparation of palmitate-BSA and oleate-BSA conjugates

To prepare a 6 mM 6:1 fatty acid-BSA conjugate solution (i.e. 6 mM fatty acid bound to 1 mM BSA), 76.93 mg palmitic acid or 84.74 mg oleic acid was weighed in a clean glass tube and converted to sodium salt by adding 200 µL 10 M NaOH and 5 mL 90°C H2O. The mixture was incubated at 90°C water bath for 10 minutes with intermittent mixing to dissolve. In parallel, 3.3 g fatty acid-free, low endotoxin BSA (Sigma A8806) was dissolved by adding 25 mL H2O, 5 mL 10X base HBSS (1078.03 mM NaCl, 45.61 mM KCl, 12.65 mM CaCl2, 4.49 mM NaH2PO4, 4.06 mM MgSO4, 3.94 mM MgCl2, 3.36 mM Na2HPO4), 1.2 mL 0.89 M NaCl, 1 mL HEPES-NaOH (pH 7.4) on a 50°C hot plate with stirring. The fatty acid solution (kept in 90°C water bath) were then added slowly and dropwise using a glass Pasteur pipette (prewarmed to 90°C) to the BSA solution rapidly stirred on a 50°C hot plate. The resulting fatty acid-BSA conjugate solution was clear without visible precipitates. After titrating the pH of the fatty acid-BSA conjugate solution to 7.4 with 1 M HCl, the total volume was brought to 50 mL with H2O in a volumetric flask. The fatty acid-BSA conjugate was filtered through a 0.22 µm syringe filter (pre-cleaned with H2O to remove wetting agents), aliquoted, and stored at -20°C. Stock solutions (6 mM; FA:BSA = 6:1) of [U-^13^C16]Palmitate-BSA conjugate and [U-^13^C18]oleate-BSA conjugate were prepared in the same way.

### 4.7 Preparation of liver co-culture medium (L-CCM) and MASH medium

L-CCM is a chemically defined, serum-free medium designed in-house for the co-culture of HepOrgs, KCs, and HSCs. CCM4 was prepared by supplementing the following components to phenol-red free William’s medium E: 2 mM Glutamax, 50 µg/mL Normocin, 0.5X MEM amino acids solution (Gibco 11130051), 1X MEM non-essential amino acids solution (Gibco 11140050), 0.1% (w/v) Poloxamer 188, lipid supplement (final concentration: 0.02 µg/L arachidonic acid, 1 µg/L linoleic acid, 1 µg/L linolenic acid, 0.05 µg/L (±)-α-lipoic acid, 0.7 µg/L (±)-α-tocopherol, conjugated to 0.1% [w/v] fatty acid-free BSA), 2.5 ng/mL murine heparin-binding EGF-like growth factor (HB-EGF), 2.5 ng/mL murine epidermal growth factor (EGF), 5 ng/mL hepatocyte growth factor (HGF), and 5 ng/mL murine macrophage colony-stimulating factor (M-CSF), 0.5 µg/mL insulin (from bovine pancreas), 0.1 µg/mL glucagon (synthetic), 10 µg/mL human transferrin (partially saturated), 2 mM sodium lactate, 50 µM choline, 0.1 mM galactose, 0.1 mM mannose, 0.1 mM taurine, 0.1 mM L-carnitine, 20 µM ethanolamine, 20 µM putrescine, 20 µM hypoxanthine, 1 µM thymidine, 0.1 µM A83-01, 20 nM selenite, 0.2 nM triiodothyronine, and 2 ng/mL dexamethasone. A83-01-supplemented L-CCM is used within 1 week.

To prepare the MASH medium (400 µM palmitate-BSA conjugate, 200 µM oleate-BSA conjugate, 100 µM cholesterol-methyl-β-cyclodextrin [MβCD] in CCM4), 4 µmole cholesterol (complexed to MβCD at 5:1 ratio; Sigma C4951) was weighed in a 50 mL tube. The cholesterol-MβCD powder was dissolved by adding 2.667 mL warm (37°C) 6 mM palmitate-BSA conjugate (6:1) and 1.333 mL warm (37°C) 6 mM oleate-BSA conjugate (6:1). The mixture was incubated in a 37°C water bath with occasional mixing. As soon as the cholesterol-MβCD powder is dissolved, the total volume was brought to 40 mL with warm L-CCM (37°C) and sterile filtered through a 0.22 µm syringe filter unit (pre-cleaned with H2O to remove wetting agents). Because fatty acid-free BSA extracts fatty acids and lysophospholipids from cells and binds hydrophobic hormones with its exposed hydrophobic binding sites^72^, 1X modified HBSS (in which fatty acid-BSA conjugates were dissolved) was used as the control to avoid extraction artefacts of fatty acid-free BSA. The control medium was prepared by mixing 4 mL 1X modified HBSS (129.19 mM NaCl, 4.56 mM KCl, 1.27 mM CaCl2, 0.45 mM NaH2PO4, 0.41 mM MgSO4, 0.39 mM MgCl2, 0.34 mM Na2HPO4, 20 mM HEPES-NaOH, pH 7.4) and 36 mL warm (37°C) L-CCM and sterile filtered through a 0.22 µm syringe filter unit.

### 4.8 Assembly of MASH-in-a-dish co-culture system and MASH induction

To assemble the MASH-in-a-dish, the culture of murine HepOrgs, primary murine KCs, and murine HSCs were synchronized. The HepOrgs were grown to 30-50 µm in diameter in expansion medium and then switch to L-CCM for maturation for 3 days. The matrix of HepOrg were then removed by dispase II treatment without dissociation of HepOrgs. After thorough removal of dispase II by washing with PBS supplemented with 10% (v/v) FBS, the HepOrgs were suspended in basement membrane extract (BME) matrix at a density equivalent to about 5 × 10^5^ cells/mL. To plate the HepOrgs on the underside of the Transwell membrane, the 6.5 mm polyester Transwell insert with 0.4 µm pores (Corning 3470) was inverted, sitting on top of the lid of a 24-well plate. Ten microliter of HepOrg BME suspension was carefully pipetted onto the centre of the underside of the inverted Transwell and allowed to sit for 2 minutes at room temperature before carefully place back the bottom 24-well plate. The whole plate with Transwell inserts inside was then inverted back and incubate in a 5% CO2, 37°C incubator for 30 minutes to gel. Meanwhile, the murine HSC were detached and resuspended at 860,000 cells/mL in L-CCM. After the BME matrix has set, 150 µL murine HSC cell suspension (about 1.29 × 10^5^ cells / insert) was added to the insert and 650 µL L-CCM was added to the bottom. Primary murine KCs were seeded at about 1.96 × 10^5^ cells / well in a separate TC-treated 24-well plate in KCM. When primary KCs were replaced by the immortalized KC cell line KUP5, KUP5 cells were seeded at 1.6 x 10^4^ cells / well because they do not stop proliferating in MASH-in-a-dish. All cells were allowed to settle overnight. The next day, the murine KCs were refreshed with 650 µL CCM4. The Transwell insert, containing murine HSC in the insert and a BME droplet of hepatocyte organoids underneath, was refreshed with 150 µL CCM4 and transferred to the KC plate to complete the assembly of MASH-in-a-dish. When examining the interaction between HepOrg and the non-parenchymal cell (NPC, KCs and HSCs), HepOrg, KCs, or HSCs were omitted. When HepOrg was omitted, an BME gel was plated as a control. HepOrg, KCs, and HSCs were co-cultured first for 3 days for adaptation. For MASH induction, the cells were treated with MASH medium (L-CCM supplemented with 400 µM palmitate-BSA conjugate [6:1], 200 µM oleate-BSA conjugate [6:1], and 100 µM Cholesterol–methyl-β-cyclodextrin complex [5:1]) for 72-96 hours.

### 4.9 Liquid chromatography-mass spectrometry (LC-MS)-based metabolomics and [U-^13^C6]-glucose stable isotope tracing in MASH-in-a-dish

After 3-day MASH or control treatment, the media of MASH-in-a-dish were refreshed with respective MASH or control media where glucose was replaced by the same concentration of ^13^C6-glucose (10 mM) for stable isotope tracing. The cells were incubated with ^13^C6-glucose for 24 hours. For metabolomics analysis of the medium, 10 µL medium from the well was harvested and mixed with ice-cold 990 µL metabolomics extraction solution (MES, methanol : acetonitrile : H2O = 2 : 2 : 1 [v/v/v]). For metabolomics analysis of the cells, the medium in the well and in the insert were removed and all cells, including the hanging HepOrg droplet, were washed twice with ice-cold PBS. The KCs were lysed in 250 µL MES. The BME droplet (HepOrg) was carefully scraped off the underside of the Transwell membrane with ethanol-cleaned spatula and transferred to an Eppendorf tube with 0.5 mL ice-cold MES. The underside of the Transwell membrane was wiped with cotton gauze to completely remove residual HepOrg cells and BME gel. Then the HSC in the Transwell insert was lysed in 250 µL ice-cold MES. All medium and cell metabolomics samples were centrifuged at 20,000 x *g* for 10 minutes at 4 °C to remove cell debris and protein precipitates. The supernatants were collected for LC-MS analysis.

LC-MS analysis was performed on an Exactive mass spectrometer (Thermo Scientific) coupled to a Dionex Ultimate 3000 autosampler and pump (Thermo Scientific) or Q-Exactive HF mass spectrometer (Thermo Scientific) coupled to a Vanquish autosampler and pump (Thermo Scientific) as previously described^73^. The MS operated in the polarity-switching mode with spray voltages of 4.5 kV and -3.5 kV. Metabolites were separated using a Sequant ZIC-pHILIC column (2.1 x 150 mm, 5 μm, guard column 2.1 x 20 mm, 5 μm; Merck) with elution buffers acetonitrile (A) and eluent B (20 mM (NH4)2CO3, 0.1% NH4OH in ULC/MS grade water (Biosolve)). Gradient ran from 20% eluent B to 60% eluent B in 20 minutes, followed by a wash step at 80% and equilibration at 20%. Flow rate was set at 150 μL/min. Analysis was performed using TraceFinder software (Thermo Scientific). Metabolites were identified and quantified on the basis of exact mass (mass tolerance ± 5 ppm) and retention time matching to standards. Peak intensities of intracellular metabolomes were normalized based on the total metabolite signal per sample. Peak intensities of extracellular (i.e. medium) metabolomes were not normalized. The isotopologue distributions were corrected for natural abundance of ^13^C. The total signal intensity of each metabolite was calculated as the sum of the signal intensities of its detected isotopologues. Data processing and analysis were performed in R (version 4.4.0). Please see Supplementary Spreadsheets 1 and 2 for signal intensities of quantified metabolites in cells and media of MASH-in-a-dish co-cultures, respectively.

### 4.10 [U-^13^C6]-Glucose and [U-^13^C16]-palmitate/[U-^13^C18]-oleate tracing in KUP5 cells

KUP5 cells were seeded at 6000 cells per well in KUP5 medium in 24-well plates and incubated overnight to attach. The medium was switched to L-CCM4 for a 24-hour adaptation. Subsequently, the cells were refreshed with the flux media: control medium with [U-^13^C6]-Glucose, control medium with [U-^13^C6]-Glucose + 100 ng/mL LPS (Invivogen, #tlrl-3pelps) + 100 ng/mL murine IFN-γ (Miltenyi Biotec, #130-105-785), L-CCM with [U-^13^C6]-Glucose and unlabelled MASH lipid mix, L-CCM with ^13^C MASH lipid mix (400 µM [U-^13^C16]-palmitate-BSA [6:1], 200 µM [U-^13^C18]-oleate-BSA [6:1], 100 µM unlabelled cholesterol-MβCD). After 24-hour incubation, the medium and KUP5 cells were extracted in metabolomics extraction solution (MES, methanol : acetonitrile : H2O = 2 : 2 : 1 [v/v/v]) for metabolomics as described in 4.9.

### 4.11 Histology and immunohistochemistry (IHC)

HepOrgs in BME (volume = 10 µL) was fixed in 650 µL 4% PFA / 5% sucrose / 50 mM NaPi (pH 7.4) for 30 minutes and then scraped and stored in a falcon tube with 10 mL 70% (v/v) ethanol at 4°C. For paraffin embedding, HepOrgs in BME was embedded in warm 2% (w/v) agarose gel and allowed to set at room temperature. Subsequently, the agarose-embedded sample was dehydrated with graded ethanol series, cleared in xylene, infiltrated with paraffin wax, and embedded in paraffin on the Slee MPS-P2 embedding centre. The paraffin block was sectioned on the Slee cut 5062 microtome at 4 µm thickness and transferred onto a microscopic slide (VWR #631-1185) and stained by haematoxylin and eosin.

For γH2AX IHC, the slides were dried overnight at 55°C, deparaffinized, and rehydrated. The antigen was retrieved by 10 mM citrate buffer (pH 6.0) in a microwave. After rinsing with PBS, endogenous peroxidases were inactivated using fresh 1% (w/v) H2O2 in methanol. The slide was subsequently washed in PBST, blocked in 10% (v/v) normal goat serum. Subsequently, the slide was incubated with primary antibody overnight in a humidity chamber at 4°C, biotinylated secondary antibody for 30 minutes at RT, and ABC/PO complex (1:25 in PBS). The signal was developed with freshly made DAB substrate for 10 minutes at RT. After rinsing away excessive substrate, the slide was counterstained with haematoxylin and rinsed in running tap water. For each experimental condition, 8 to 12 representative fields (40× magnification) per stained slide were analyzed. The number of gH2AX-stained nuclei was counted and expressed as a percentage of total nuclei using ImageJ software (NIH, Bethesda, MD, USA) with the Multipoint tool.

The histological features of MASH (previously NASH) are typically scored according to Kleiner et al.^74^ Whilst not all elements of this scoring system are present in the MASH-in-a-dish model, the degree of steatosis and incidence of ballooning was assessed according to the criteria outlined by Kleiner et al. Steatosis was reported based on the presence of multiple clearly defined cytoplasmic vacuoles. Samples were attributed a score of 0 (none), 1 (some) or 2 (all) according to the proportion of cells showing steatosis. Ballooning is defined by hepatocellular hypertrophy to 1.5×-2× the size of adjacent hepatocytes, cytoplasmic rarefaction and a central nucleus and was scored 0 = absent or 1 = present. A score of 0.5 was assigned to uncertain cases.

Histological scoring was performed by a pathologist blinded to treatment conditions using coded samples. The treatment codes were revealed only after completion of the scoring. Images were acquired by an independent researcher. Please refer to Supplementary Table 1 for antibody source and dilution.

### 4.12 Immunofluorescence (IF)

Cells were seeded on coverslips and fixed in 4% (w/v) paraformaldehyde / 5% (w/v) sucrose / 50 mM NaPi buffer, pH 7.4, for 15 minutes at room temperature. After fixation, the cells were washed twice with PBS and permeabilized in 1% (w/v) Triton X-100 in TBST (150 mM NaCl, 20 mM Tris-HCl, 0.05% [w/v] Tween 20, pH 7.6) for 10 minutes and washed twice with TBST. The coverslips were then incubated with unconjugated primary antibodies diluted in TBST for 1 hour. After washing 3 times with TBST, cells were incubated with secondary antibodies in TBST for 1 hour. After washing 3 times with TBST, the coverslips were mounted in Vectashield with DAPI (Vector laboratory, #H-1200-10). Images were taken with a Leica DM6000 microscope using LAS AF software (Leica).

Please refer to Supplementary Table 1 for antibody source and dilution.

### 4.13 Neutral lipid staining and confocal microscopy

After co-culture and treatment with control medium or MASH medium, HepOrgs in BME was fixed in 4% (w/v) paraformaldehyde / 5% (w/v) sucrose / 50 mM NaPi (pH 7.4) / 0.05% (w/v) glutaraldehyde for 30 minutes at room temperature. To visualize lipid droplets, HepOrgs were stained with Bodipy 493/503 or Lipidspot610. For Bodipy 493/503 staining, cells were incubated with 0.25 µg/mL Bodipy 493/503 in TBST for 5 minutes at room temperature and then wash 3 times with 3 TBST. For LipidSpot610 staining, HepOrgs were incubated with 1:1000 LipidSpot610 in staining buffer (5 mM Na2SO4 and 5 mM EDTA-NaOH [pH 8.0]) for at least 4 hours at room temperature. Cells stained with LipidSpot610 were imaged in staining buffer with LipidSpot610 without washing.

Fixed Transwell samples were placed into a 3510 FluoDish (WPI, UK) and stacks (step size 0.5936 um with a range to cover about ¾ of the larger organoids) were imaged using a Leica Stellaris WLL Falcon system with a 25× water immersion Objective (HC Fluotar L VISIR, NA0.95), zoomed 3.18×, in bidirectional mode. Resulting images had 86.4 nm pixel size and were 1688 pixels wide and high. Ilumination was performed at 405 nm (1.5% laser power diode laser illumination for Hoechst 33342 emission range 415-553 nm) and 610 nm (0.85% WLL laser power for LipidSpot610 and emission range 616-726 nm) with HyD-S and HyD-X in analogue/digital more, averaging 3×. Voxel dimensions were 0.264 × 0.264 × 0.5936 µm^3^ and a 0.25 µs pixel dwell time. Bright field images were simultaneously collected by the transmission detector. Images were denoised and deconvolved using the Lightning confocal mode in the LAS-X software. The Hoechst 33342 staining was less effective in the organoids that accumulated large lipid droplets, therefore the nuclear image intensity in these samples were adjusted differently to show the weakly labelled nuclei slightly better. All other channels intensity levels were adjusted equally. Maximum intensity projection (MIP) images are shown in Figure 1B. Three-dimensional maximum intensity projection (3D-MIP) video renderings of selected HepOrgs are shown in the Supplementary Videos 1-3.

Spinning disk confocal imaging was performed on an Evident SpinSR10 system equipped with a Yokogawa W1-SoRa spinning disk mounted on a IX83 stand with Hamamatsu ORCA Flash 4.0 cameras (Evident, Leidendorp, the Netherlands). The system was run by Evident cellSens Dimension software (vs 4.2) in 50-µm pinhole disk confocal mode using a 20× LUCPLFLN air objective (NA 0.45, rendering a 0.325 µm pixel size). Samples were clamped in a custom-designed and in-house fabricated Transwell holder, which positioned the outer filter surface in close proximity to the coverslip (STL/OBJ files available upon request). Z-stacks were acquired at 1.5 µm intervals, with the imaging range covering the entire organoid volume. Neutral Lipids (Bodipy 493/503) were illuminated with the 488 nm laser line (16 mW power), while Hoechst33342 channels were detected using the 405 nm line (16.5 mW power), exposed for 100 ms and 200 ms, respectively. A quad band dichroic was used and emission filters were 525/50 nm and 447/60 nm respectively. Maximum projections of cropped regions with representative organoids are shown (Figure S1M).

Microscopy for 2D HepOrg on collagen-coated surface (Figure S4B) was performed on a Nikon A1R confocal microscope (Nikon Europe) with a 40x ELWD air objective (NA 0.4). Single Z planes were collected using the 405 and 645 nm of the LUA4 laser set run at 2.2 mW and 1.5 mW laser powers, passed through a Quad Band (405/488/561/640) dichroic with emission filters 450/50 and 700/75, for the Hoechst33342 and Lipidspot610 channels respectively. Pinhole was set to 1.75 AU (640nm as reference), 1x zoom in 512x125 scan mode resulting in a pixel size of 0.62 um. Tile scans of 4x4 images were taken per well, with an autofocus routine to find optimal nuclei contrast for each well position. Images were denoised using the ai module in NIS elements (version 5.1) and uniformly intensity adjusted.

### 4.14 Measurement of reactive oxygen species (ROS) production rate by dihydroxyethidium (DHE)

The ROS production rate was measured by DHE oxidation in a live cell assay in a CLARIOstar fluorometer (BMG) at 37°C in ambient atmosphere. Reaction of DHE with ROS yields 2-hydroxyethidium and ethidium, the fluorescence of both can be monitored at λEx/λEm = 520-20/590-40. HepOrgs were seeded on 96-well plates coated with 5 µg/cm^2^ type I collagen from rat tail. Cells were treated 4 days with MASH or control CCM. To quantify ROS production, cells were washed once with 100 µL assay medium (minimum essential medium supplemented with 21 mM NaCl, 20 mM HEPES-NaOH, 2 mM glutamine, pH 7.4) and incubated in 100 µL assay medium. After monitoring the background fluorescence for 1 hour, 10 µM DHE (final concentration; abcam #ab145360) was added to monitor ROS production rate. The ROS production rate was calculated from the slope of the fluorescence kinetic curve, corrected for background, and then normalized to cell numbers per well.

### 4.15 Double-stranded DNA (dsDNA) microplate assay

To quantified cell numbers, KCs and HSCs in microplates were fixed for 15 minutes in 4% (w/v) paraformaldehyde (PFA) / 5% (w/v) sucrose / 50 mM Na2HPO4-NaH2PO4 (pH 7.4). After removing the fixative, the cells were washed twice with assay buffer (5 mM Na2SO4 and 5 mM EDTA-NaOH [pH 8.0]) and stored in assay buffer at 4°C until dsDNA staining. For hepatocyte organoids culture in basement membrane extract (BME), cells were fixed in ice-cold metabolomics extraction solution (MES methanol : acetonitrile : H2O = 2 : 2 : 1 [v/v/v]) for 30 minutes with 1 refreshment after the first 15 minutes. Because BME contains dsDNA, an empty BME control was included for each assay timepoint to control for the background dsDNA signals. After aspirating MES, the HepOrg-containing BME dome was dried to a thin disk at 4°C overnight. dsDNA of cells were stained with 5 µg/mL Hoechst33342 (Sigma) in assay buffer for at least 24 hours to allow stoichiometric binding between dsDNA and Hoechst33342. The fluorescence of the microplate were then read in CLARIOstar fluorometer at λEx/λEm = 350-10/490-40, room temperature. A dsDNA standard curve of highly polymerized calf thymic DNA from 20 µg/mL to 0.3125 µg/mL (prepared 5 µg/mL Hoechst33342 in assay buffer) was measured in the same plate for dsDNA quantification. The interpolated dsDNA content was then converted to cell numbers based on the cellular dsDNA content of a normal diploid murine cell (6 pg per diploid murine cell).

### 4.16 Quantification of LipidSpot610 neutral lipid staining in microplates by fluorometer

Hepatocyte organoids were seeded in 48-well plates coated with rat tail collagen, Type I, 5 µg/cm^2^. After treatment, cells were fixed in 100 µL 4% (w/v) PFA / 5% (w/v) sucrose / 50 mM NaPi (pH 7.4) per well for 15 minutes at room temperature. The fixative was removed and the fixed cells washed 2 times with staining buffer (5 mM Na2SO4 and 5 mM EDTA-NaOH [pH 8.0]). Cells were stained with 200 µL LipidSpot610 (Biotium) working solution (1:1000 dilution in staining buffer) for at least 4 hours. The signal of LipidSpot610 quantified by well scan in a CLARIOstar fluorometer at λEx/λEm = 600-20/690-100. Because LipidSpot610 is a fluorogenic environment-sensitive lipid probe, the measurement was performed without removing excessive LipidSpot610 to avoid re-partition of LipidSpot610 out of lipid droplets. The LipidSpot 610 signal was normalized to dsDNA content per well.

### 4.17 Sandwich enzyme-linked immunosorbent assay (sELISA)

sELISA for IL-6 (R&D, #DY-406), TNF-α (R&D, #DY-410), TGF-β1 (R&D, #DY-1679), pro-collagen 1A1 (abcam, #ab216791), and ApoB (abcam, #ab230932) were performed according the instruction of the manufacturers.

sELISA for the murine albumin was set up using capture and biotinylated detection antibodies specific for the murine albumin. All steps were performed at room temperature. Briefly, in a 96-well high-binding strip plates, each well was coated with 100 µL 3 µg/mL goat anti-mouse albumin polyclonal antibody (Bethyl laboratory, #A90-134A) in PBS overnight. Unbound antibodies were washed away with PBST (PBS with 0.05% Tween 20) and the well was blocked with 300 µL 1% (w/v) BSA (Sigma, #A7030) / PBS for 1 hour. To each well, 100 µL medium sample (diluted in 1% BSA) or 100 µL mouse albumin standards (2000, 1000, 500, 250, 125, 62.5, 31.25 pg/mL in 1% BSA/PBS) were added and incubated for 1.5 hours. After 3 washings with 300 µL PBST, 100 µL 0.1 µg/mL biotinylated anti-mouse albumin polyclonal antibody in 1% BSA/PBS was added to each well. After 1-hour incubation, 100 µL Streptavidin-HRP (horseradish peroxidase, R&D #DY-998; 1:40 dilution in 1% BSA/PBS) was added and the plate was further incubated for 15 minutes. After 3 washings with PBST, the signal was developed with 100 µL TMB ELISA substrate (R&D, #DY998B). The reaction was stopped by adding 50 µL 1 M H2SO4 and the absorbance was measured at 490 nm.

Concentrations of target proteins in the medium samples were calculated by interpolating the sample absorbance with a standard curve generated by fitting the standards to the 4-parameter logistic regression model.

### 4.18 Sodium dodecyl sulfate–polyacrylamide gel electrophoresis (SDS-PAGE) and western blot

After treatment, whole cell lysates were prepared in RIPA buffer (Radioimmunoprecipitation assay buffer: 150 mM NaCl, 1% [w/v] Nonidet P-40, 0.5% [w/v] Na deoxycholate, 0.1% [w/v] SDS, 50 mM Tris-HCl [pH 8.0]) supplemented with 5 mM EDTA-NaOH, 1X Halt™ Protease Inhibitor Cocktail (Thermo Scientific #78437) and 1X PhosSTOP phosphatase inhibitor cocktail (Sigma #4906845001). Protein content was quantified by Pierce™ BCA Protein Assay Kits (Thermo Scientific #23227). Equal amounts of protein (20 to 40 μg per lane) were subjected to reduced SDS-PAGE, transferred to 0.45 µm PVDF membranes (Sigma, #IPVH00010) by semi-dry blotting and blocked overnight in 5% (w/v) fatty acid-free BSA / TBST at 4°C. For immunodetection, the PVDF membranes were incubated with the primary antibody for 1 hour, followed by 3 washes with TBST, 1 hour incubation with horseradish peroxidase-conjugated secondary antibody, and 3 washes with TBST. All antibodies were diluted in 1% (w/v) fatty acid-free BSA / TBST and the antibody incubation was performed at room temperature. The PVDF membrane was developed with SuperSignal™ West Pico PLUS Chemiluminescent Substrate (Thermo Scientific, #34577) and detected by ChemiDoc imager (Bio-Rad). Densitometry was performed using ImageJ.

Please refer to Supplementary Table 2 for antibodies and dilutions.

### 4.19 Enzymatic activity assays for aspartate transaminase (AST) and L-lactate dehydrogenase (LDH)

AST activity in culture medium was determined using a commercial kit (IFCC mod. liquiUV, HUMAN Diagnostic, 12011) with modification for 96-well microplates (Greiner Bio-One UV-Star 96-well UV spectroscopy plates, #675801). A master mixture was prepared by mixing 4 parts BUF (100 mM Tris buffer (pH 7.9), 300 mM L-aspartate, ≥ 1.13 U/mL LDH, ≥ 0.75 U/mL MDH, 0.095% sodium azide) with 1 part SUB (60 mM 2-oxoglutarate, 0.9 mM NADH, 0.095% sodium azide0). Both the master mix and the microplate were pre-warmed to 37°C. In a 96-well microplate, 40 µL spent culture medium was added, followed by 200 µL master mix. The plate was then immediately read in a spectrophotometer with absorbance at 340 nm (A340) recorded every 2 minutes with pathlength correction. The slope of A340 was used to calculated AST activity in the sample and expressed as U/L. One unit of AST activity is defined as 1 µmol NADH oxidation per minute at assay condition at 37°C.

Medium LDH activity was measured by a spectrophotometric assay in the forward reaction. Briefly, 20 µL medium was transferred into a pre-warmed UV-Star 96-well plate. Subsequently, 200 µL assay master mix (0.1 M K2HPO4-KH2PO4 [pH 7.4], 0.2 mg/mL NADH, and 10 mM Na-Pyruvate) was added and the measurement of absorbance at 340 nm was immediately started and measured nm every 2 minutes with pathlength correction. The slope of A340 was used to calculated LDH activity in the sample and expressed as U/L. One unit of LDH activity is defined as 1 µmol NADH oxidation per minute at assay condition at 37°C.

### 4.20 Bile salt measurement by HPLC

Bile salt species were separated and quantified by reverse-phase HPLC, which was an adaptation to the method used by Kunne et al^75^. To evaluate bile salt secretion of HepOrgs, 100 µL 3-day HepOrg-conditioned CCM or fresh CCM were deproteinized by addition of 500 µL acetonitrile. Sample were incubated on ice for 1 hour for deproteinization. Following centrifugation (10 min, 4°C, 14000 x *g*), the supernatant was harvested. The solvent was evaporated from the supernatant in a SpeedVac Vacuum Concentrator. The pellet was solubilized in 200 µL 25% (v/v) methanol. One hundred microliter sample was applied to a Hypersil C18 HPLC column (internal diameter: 3 µm, column length: 15 cm; Thermo Scientific, Breda, The Netherlands) operated at 20°C. The starting eluent consisted of 6.8 mM ammonium formate (pH 4.2), followed by linear gradient or isocratic elution with acetonitrile at the indicated concentration: 0-27% (0-1 min), 27% (1-4 min), 27-29% (4-10 min), 29-32% (10-17 min), 32% (17-23 min), 32-36% (23-27 min), 36-45% (27-31 min), 45-52% (31-36 min), 52-76% (36-46 min), 76-100% (46-48 min), 100% (48-50 min), 100-0% (50-52 min), and 0% (52-58 min). The flow rate was 0.8 mL per minute. Detection was performed using a CAD (Charged Aerosol Detector). Quantification of the different bile salt species was performed by using a calibration curve for all different bile salt species.

### 4.21 Triglyceride assay

To quantify triglyceride in HepOrgs, a sensitive fluorescent triglyceride microplate assay was developed. Triglyceride was quantified through the NADH generated by the coupled reactions of lipoprotein lipase and glycerol dehydrogenase and validated using triolein, tripalmitin, tristearin, and trimyristin. Following treatment, the BME matrix was removed using dispase II and the HepOrgs were washed two times in PBS. The cell pellet was solubilized in ice-cold Triton X-100 lysis buffer (2% [w/v] Triton X-100, 50 mM Tris-HCl pH 7.6, 100 mM KCl, 5 mM EDTA-NaOH pH 8.0) by mixing at the maximal setting on a vortex for 5 minutes. Insoluble materials were pelleted by centrifuging at 20,000 x *g for* 5 minutes at 4°C. The supernatant was harvested for triglyceride measurement. The triglyceride assay was performed in a solid-black 96-well microplate in a CLARIOstar fluorimeter at 37°C. Throughout the assay, NADH fluorescence was monitored at λex/λem= 340-10/460-20. Briefly, 20 µL sample was mixed with 150 µL reagent A (2 mg/mL NAD^+^ and 0.13% [w/v] Triton X-100 in assay buffer; assay buffer: 50 mM ammonium sulfate – KOH [pH 10.2], 50 mM K2CO3-KHCO3 buffer [pH 10.2], 0.05% [w/v] sodium azide) in the well. Subsequently, 20 µL Reagent B (15 U/mL glycerol dehydrogenase in assay buffer) was added to detect and remove endogenous glycerol. Finally, 10 µL Reagent C (30 U/mL lipoprotein lipase in assay buffer) was added to release the glycerol backbone from triglyceride for quantification by glycerol dehydrogenase. Triglyceride in samples was quantified using triolein standards. The assay was linear between 39.1 picomole and 1250 picomole triolein.

### 4.22 Lentivirus production and transduction for shRNA-mediated knockdown and CRISPR-Cas9-mediated knockout

Lentivirus was produced as previously described with minor modification^76^. Briefly, HEK293T cells were seeded on 145 mm TC-treated culture dish in DMEM supplemented with 10%(w/v) FBS, 2 mM L-glutamine, and 100 U/L penicillin and 100 µg/mL streptomycin. At 70-80% confluency, the cells were transfected with a mixture (per 145 mm dish) of 7 µg pVSV-G, 5 µg pRSV-rev, 13 µg pMDLg/pRRE, 20 µg transfer vector of interest, and 90 µL polyethylenimine (PEI, 1 mg/mL; Polysciences, #24765) in serum-free DMEM. Cells were refreshed the next day. Lentivirus supernatant were harvested 24 hours after medium refreshment, centrifuged at 1000 *x g* for 10 minutes at 4°C to remove debris, and then clarified by filtering through 0.45 µm low-protein binding PVDF membrane filter (Millipore #SLHV033R). The lentivirus supernatant is either directly used for transduction or further concentrated by precipitation with Poly(ethylene glycol) 8000 (PEG 8000) and NaCl.

To precipitate lentivirus, 12 mL 4X lentivirus concentrator (40% [w/v] PEG-8000, 1.2 M NaCl, 200 mM HEPES-NaOH, pH 7.2) was added to 36 mL clarified lentiviral supernatant and the mixture was incubated for 4 hours to overnight on a rotator at 4°C. The lentiviral precipitates were centrifuged at 2000 *x g* for 1 hour at 4°C. After removing supernatant, the pellet was resuspending in 720 µL lentivirus storage buffer (10% [w/v] Lactose, 20 mM HEPES-NaOH, 20 mM MgCl2, pH 7.4), aliquoted, and stored at -80°C until use.

For shRNA-mediated *Cpt1a* knockdown, adherent KUP5 cells were transduced with lentivirus at 20% confluency with 10 µg/mL Diethylaminoethyl (DEAE)-dextran (Sigma D9885) and selected with 2.5 µg/mL puromycin.

For CRISPR-Cas9-mediated *Acod1* knockout, KUP5 cells were transduced in suspension by spinoculation. Briefly, single cell suspension of KUP5 cells equivalent to 20-30% confluency were seeded in a 12-well plate in 200 µL KUP5 medium, followed by 100 µL 50 µg/mL DEAE-dextran and 100 µL 50X concentrated lentivirus stock, and 100 µL KUP5 medium. The plate was centrifuge at 800 *x g* at 32°C for 30 minutes for spinoculation. After the spinoculation, 500 µL KUP5 medium were gently added to the cells. The medium was refreshed 24 hours after spinoculation. The transduced cells were cloned by limited dilution. To screening for knockout clones, the clonal lines were treated with MASH medium for 24 hours and medium itaconate was measured by LC-MS as described in Materials and Method **4.9**. Clones that did not produce itaconate following MASH medium treatment were further expanded and validated by western blot for ACOD1.

Please refer to Supplementary Table 3 for lentivirus packing plasmids, lentiviral vectors, and target sequences of shRNA and sgRNA.

## 5. Acknowledgments

This work was supported by a Veni grant from the Dutch Research Council (NWO) awarded to J.C. Chang [Grant ID https://doi.org/10.61686/HBGIJ22746]. The authors would like to express their gratitude to the following colleagues for the support received during this study: Suzanne Duijst (Tytgat Institute for Liver and Intestinal Research, Amsterdam UMC) for liver perfusion, dr. Rudi de Waart (Tytgat Institute for Liver and Intestinal Research, Amsterdam UMC) for HPLC-based bile salt quantification, and Jeroen Jansen (Faculty of Veterinary Medicine, Utrecht University) for operation of mass spectrometers. We would also like to thank Estelle Hunter, Iris van Viegen and Hristo Ivanov (Utrecht University) for their technical assistance.

## 6. Supplementary Figure Legends

**Figure S1.**
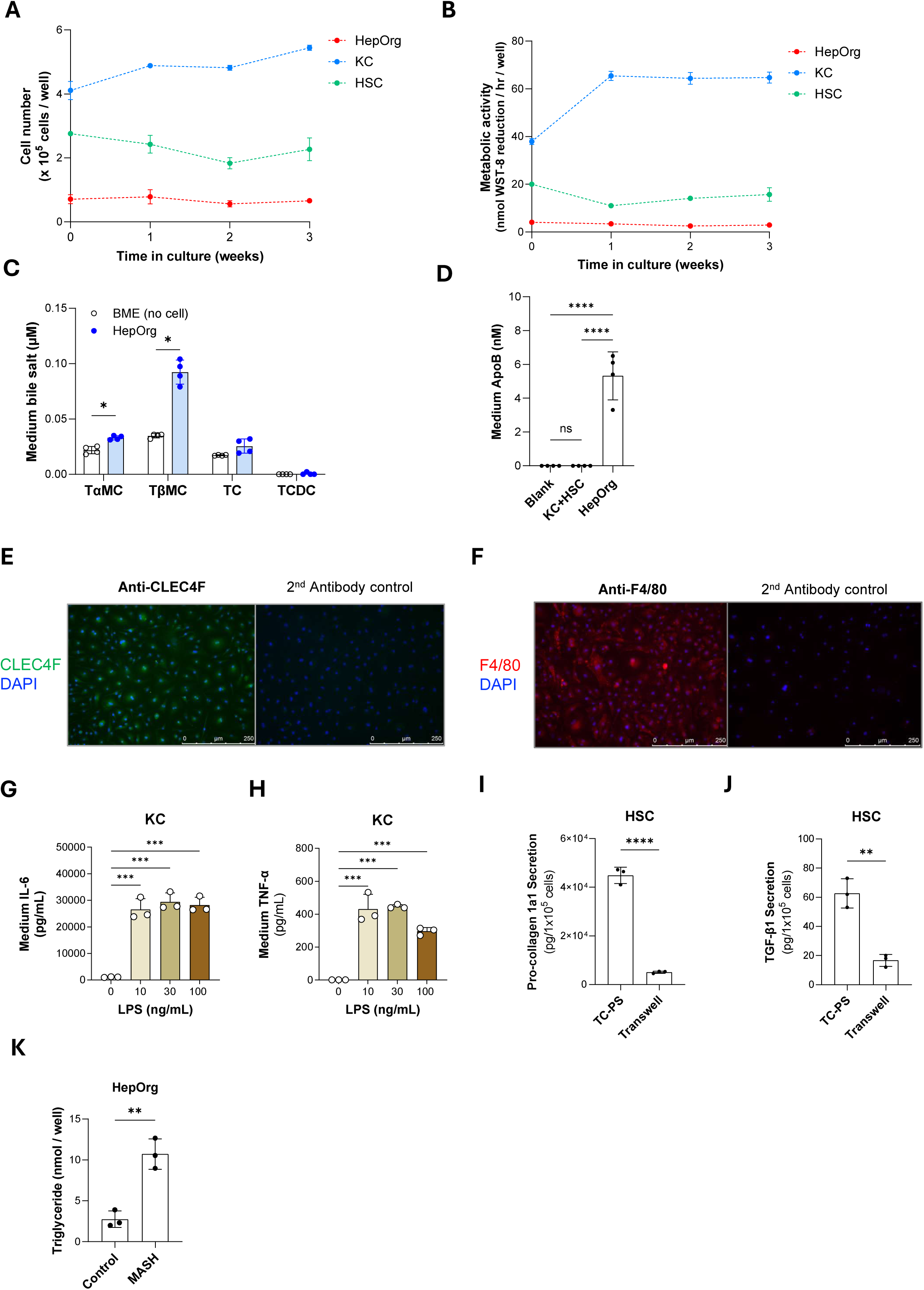
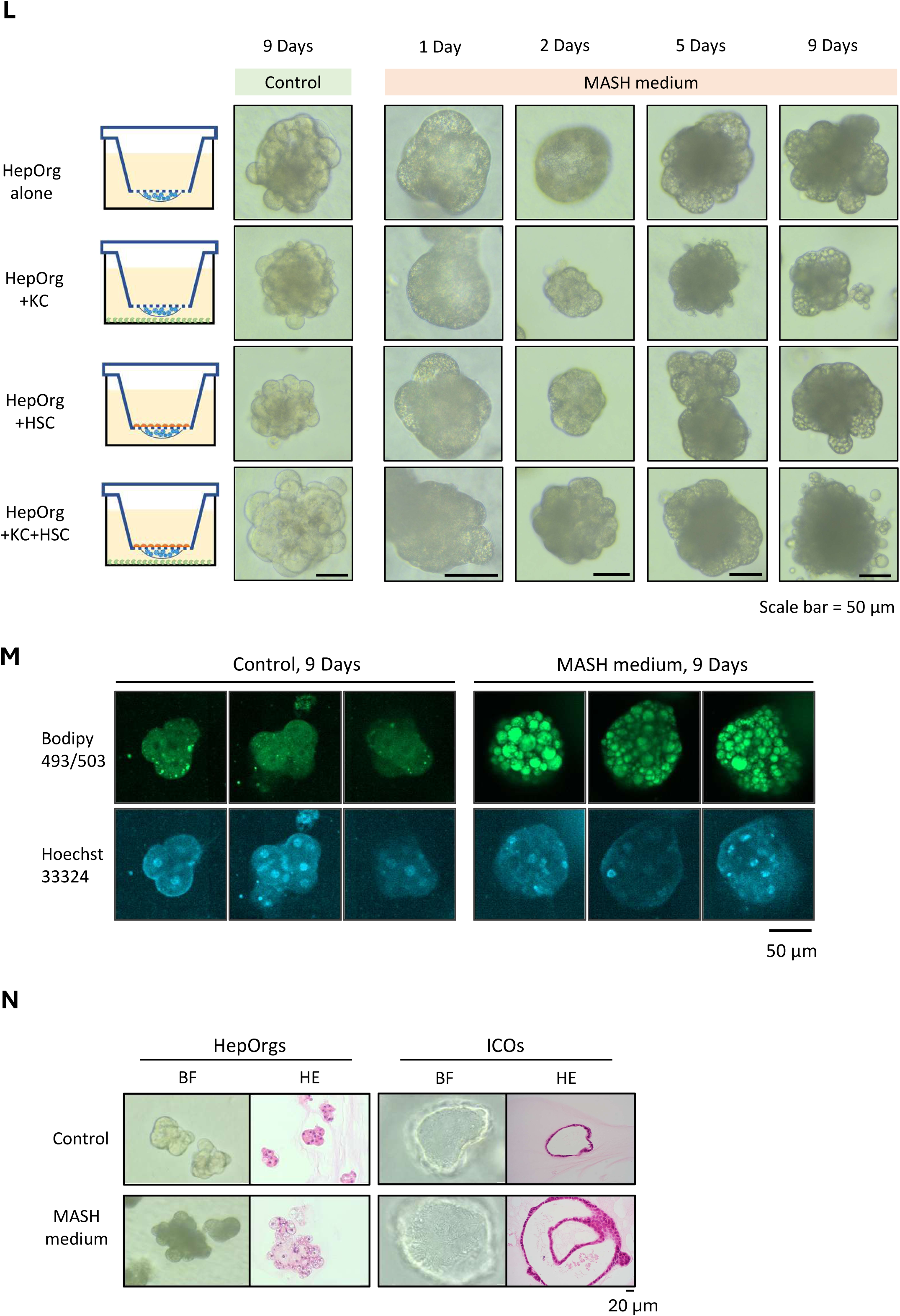
Functional characterization of murine hepatocyte organoids (HepOrgs), Kupffer cells (KCs), and self-immortalized hepatic stellate cells (HSCs). (Related to Figure 1) **(A-B)** HepOrg (cultured in BME), KCs, and HSCs were seeded in 48-well plates in their respective culture media and switched to the L-CCM for 3 weeks. Cell numbers (A) and metabolic activities (B) were assayed with dsDNA assay and WST-8 assay, respectively, at the baseline (immediately after medium switch) and after culturing in L-CCM for 1, 2, and 3 weeks. The dsDNA content in HepOrg was corrected against BME controls. n = 4 independent wells for all conditions. **(C)** HepOrgs were differentiated (as described in Methods) and subsequently incubated 3 days in L-CCM4. Bile salts secreted into media were quantified by HPLC. TαMC, tauro-α-muricholate; TβMC, tauro-β-muricholate; TC, taurocholate; TCDC, taurochenodeoxycholate. n = 4 independent wells. **(D)** To detect apolipoprotein B (ApoB) secretion, a marker of very low-density lipoprotein (VLDL), media were conditioned by HepOrgs or KCs + HSCs for 5 days. Medium ApoB was quantified by sandwich ELISA. n = 4 independent Transwell culture. **(E)** Immunofluorescence of CLEC4F (a KC-specific marker) in primary murine KCs. **(F)** Immunofluorescence of F4/80 (a general macrophage marker) in primary murine KCs. **(G-H)** Primary murine KCs were stimulated with lipopolysaccharide (LPS) at 0, 10, 30, 100 ng/mL for 24 hours. Medium IL-6 (G) and TNF-α (H) were measured by ELISA. n = 4 independent wells, representative of 2 independent experiments. **(I-J)** Activated murine HSCs were cultured on tissue-culture-treated polystyrene surface (hard, high stiffness) or polyester Transwell membrane (soft, low stiffness) for 4 days in control medium or MASH medium. Medium pro-collagen1A1 (I) and TGF-β1 (J) were measured by ELISA. **(K)** HepOrgs were treated with control L-CCM or MASH L-CCM for 4 days. Intracellular triglyceride was quantified enzymatically. n = 3 independent wells. **(L)** HepOrgs were cultured alone or with KCs, HSCs, or KCs +HSCs in control medium or MASH medium for up to 9 days. Development of micro- and macro-vesicular steatosis were observed by brightfield microscopy. **(M)** HepOrgs were cultured with HSCs for 9 days in control or MASH medium. Neutral lipids and dsDNA were stained with Bodipy 493/503 and Hoechst33342, respectively. **(N)** Murine HepOrgs and intrahepatic cholangiocyte organoids (ICOs) were treated with control or MASH lipid mix in L-CCM for 4 days. Steatosis was assessed by brightfield microscopy and HE histology. **Statistical analysis:** (C) Multiple unpaired *t*-test with Welch correction. (D) One-way ANOVA with Tukey’s multiple comparison tests. (G-H) One-way ANOVA with Dunnett’s multiple comparison tests. (I-J) Two-tailed unpaired Student’s *t*-test. (K) Two-tailed unpaired Student’s *t*-test. ns, not significant; *, *P* < 0.0332; **, *P* < 0.0021; ***, *P* < 0.0002; ****, *P* < 0.0001.

**Figure S2.**
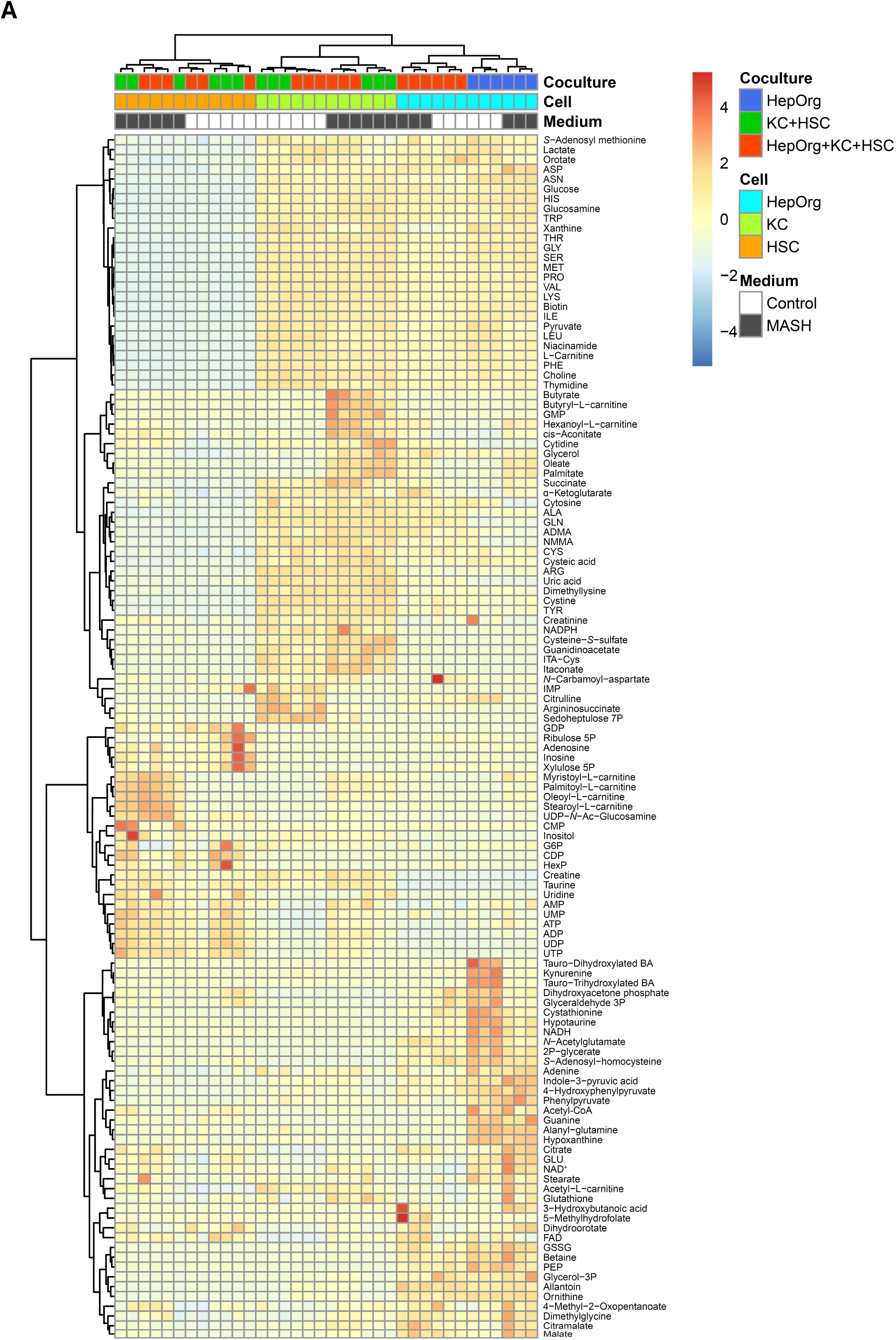
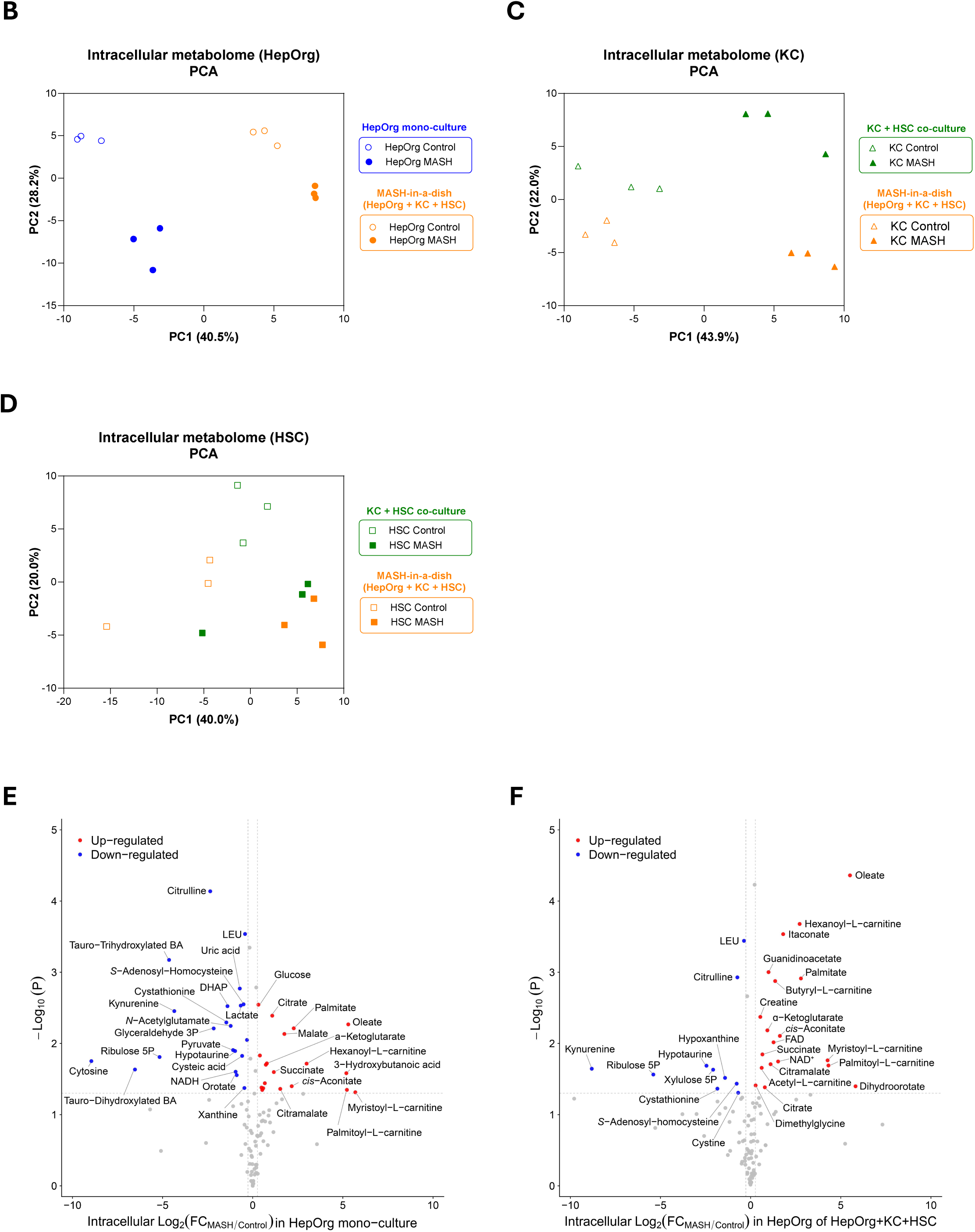
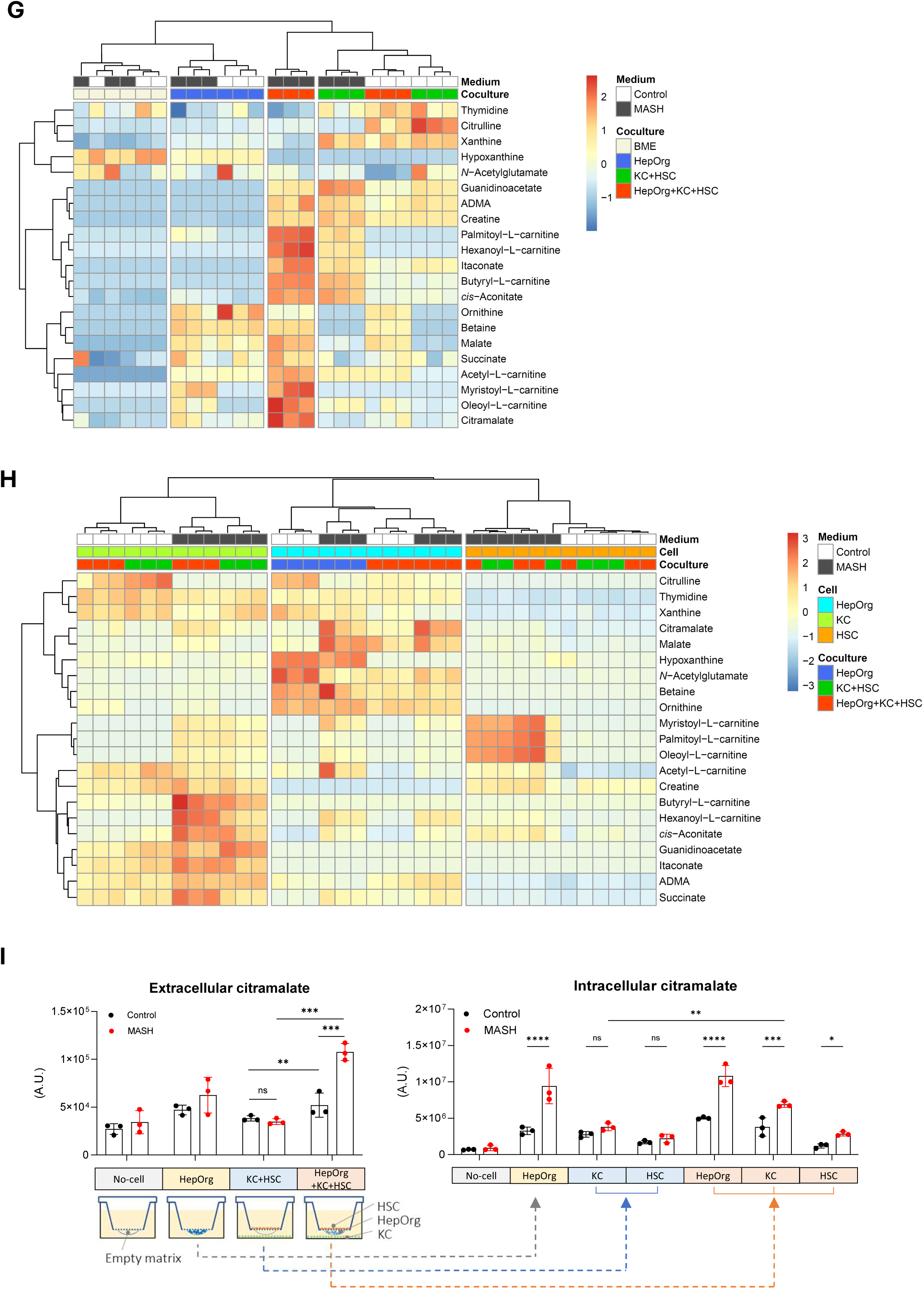
Effects of cell-cell interactions on cell-type-specific intracellular metabolomes in different control and MASH co-culture conditions. (Related to Figure 2) **(A)** Heatmap with unsupervised clustering of intracellular metabolites detected by targeted metabolomics. HepOrgs, hepatocyte organoid; KC, Kupffer cells; HSC, hepatic stellate cells. **(B-D)** Principle component analyses of HepOrg metabolomes (B), KC metabolomes (C), and HSC metabolomes (D) in different MASH-in-a-dish culture conditions. **(E)** Volcano plot of HepOrg intracellular metabolites in MASH medium versus control medium in HepOrg mono-culture. **(F)** Volcano plot of HepOrg intracellular metabolites in MASH medium versus control medium in HepOrg + KC + HSC co-culture. **(G-H)** Heatmap with unsupervised clustering of extracellular levels (G) and intracellular levels (H) of MASH-associated crosstalk metabolites under different co-culture conditions in control medium and MASH medium (see also Figure 2G). **(I)** Intracellular and extracellular (i.e. conditioned co-culture medium) levels of citramalate in different MASH-in-a-dish co-cultures; extracted from the metabolomics experiment in Figure 2A. Data presented as mean ± SD of 3 independent Transwell cultures.

**Figure S3.**
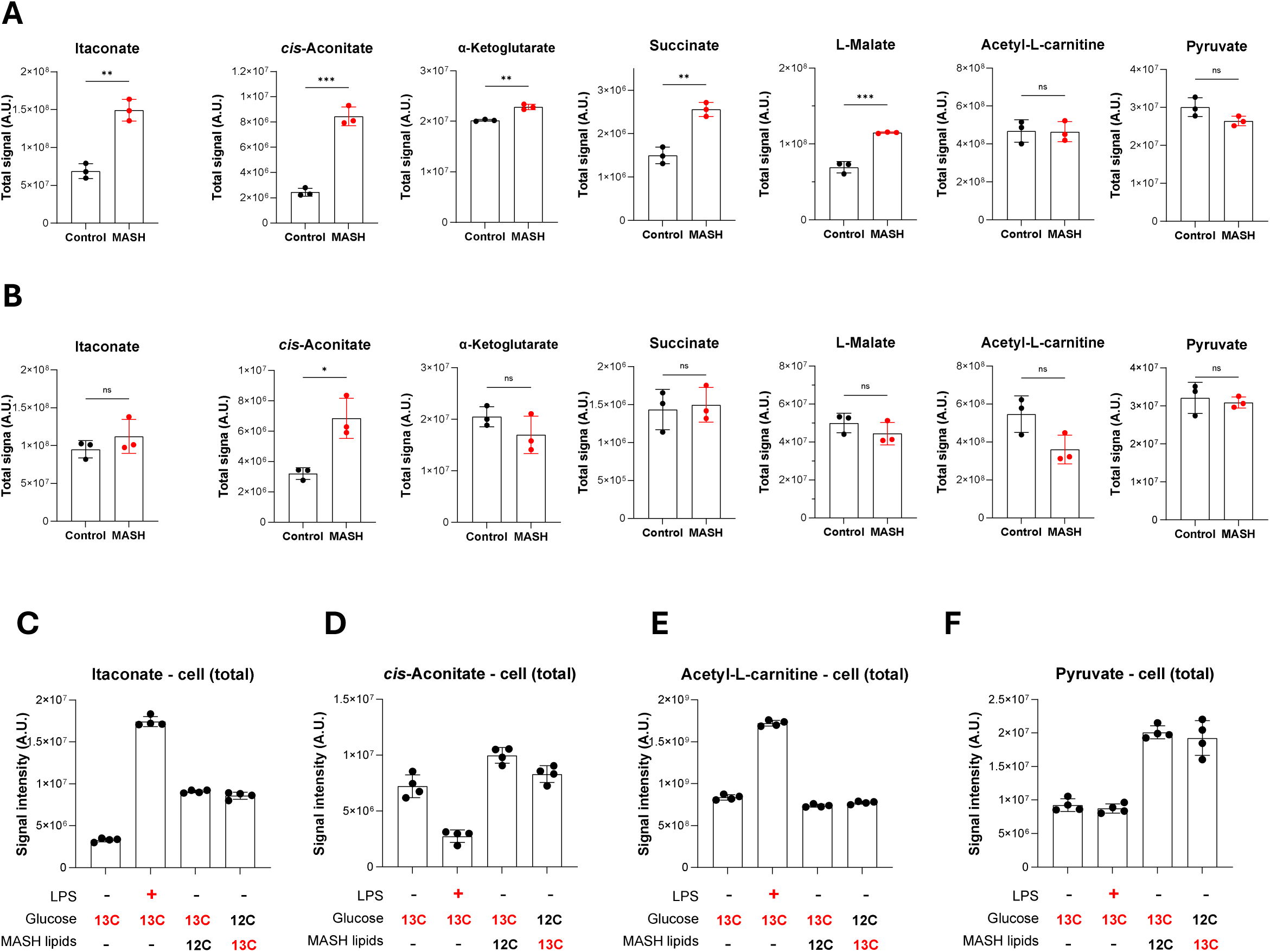
Total levels of selected intracellular metabolites in [U-^13^C6]-Glucose and [U-^13^C16]-palmitate/[U-^13^C18]-oleate tracing in KCs in the absence and presence of MASH lipid mix. (Related to Figure 3) **(A-B)** Total intracellular levels of itaconate, *cis*-aconitate, α-ketoglutarate, succinate, L-malate, acetyl-L-carnitine, and pyruvate in primary KCs of HepOrg + KC + HSC co-culture (A) and KC + HSC co-culture (B). Data derived from the experiment of Figure 2A. n = 3 independent Transwell cultures. **(C-F)** Total intracellular levels of itaconate, *cis*-aconitate, acetyl-L-carnitine, and pyruvate in KUP5 cells treated as in Figure 3K. **Statistical analysis:** (A-B) Two-tailed, unpaired Student’s *t*-test. ns, not significant; *, *P* < 0.0332; **, *P* < 0.0021; ***, *P* < 0.0002.

**Figure S4.**
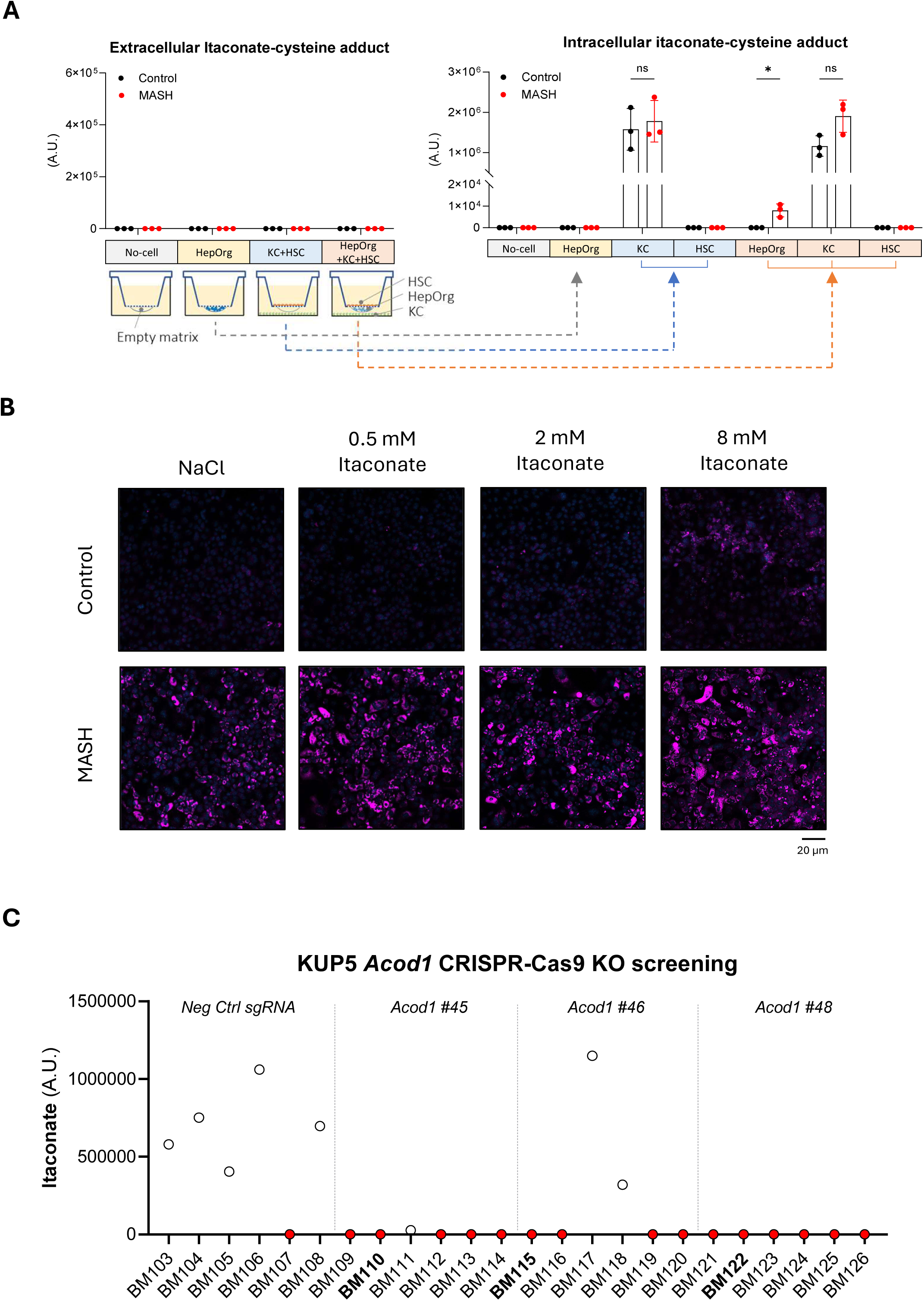
KC-derived itaconate signals to HepOrgs in MASH-in-a-dish. (Related to Figure 4) **(A)** Intracellular and extracellular (i.e. conditioned co-culture medium) levels of itaconate-cysteine adduct in different MASH-in-a-dish co-cultures; extracted from the metabolomics experiment in Figure 2A. Data presented as mean ± SD of 3 independent Transwell cultures. **(B)** Representative confocal microscopy images of Lipidspot610 neutral lipid staining of 2D HepOrgs cultured on a collagen-coated surface. The HepOrgs were treated with 0, 0.5, 2, and 8 mM itaconate in the absence or presence of the MASH lipid mix for 4 days. **(C)** After transduction with negative control sgRNA or 3 independent sgRNAs against *Acod1* (#45, #46, #48), the transduced KUP5 cells were cloned by limited dilution cloning. To screen for *Acod1* knockout clones, the KUP5 clones were treated with the MASH lipid mix for 24 hours. Itaconate levels in the media was measured by LC-MS to screen for knockout clones.

## 7. Supplementary Tables

**Supplementary Table 1.**
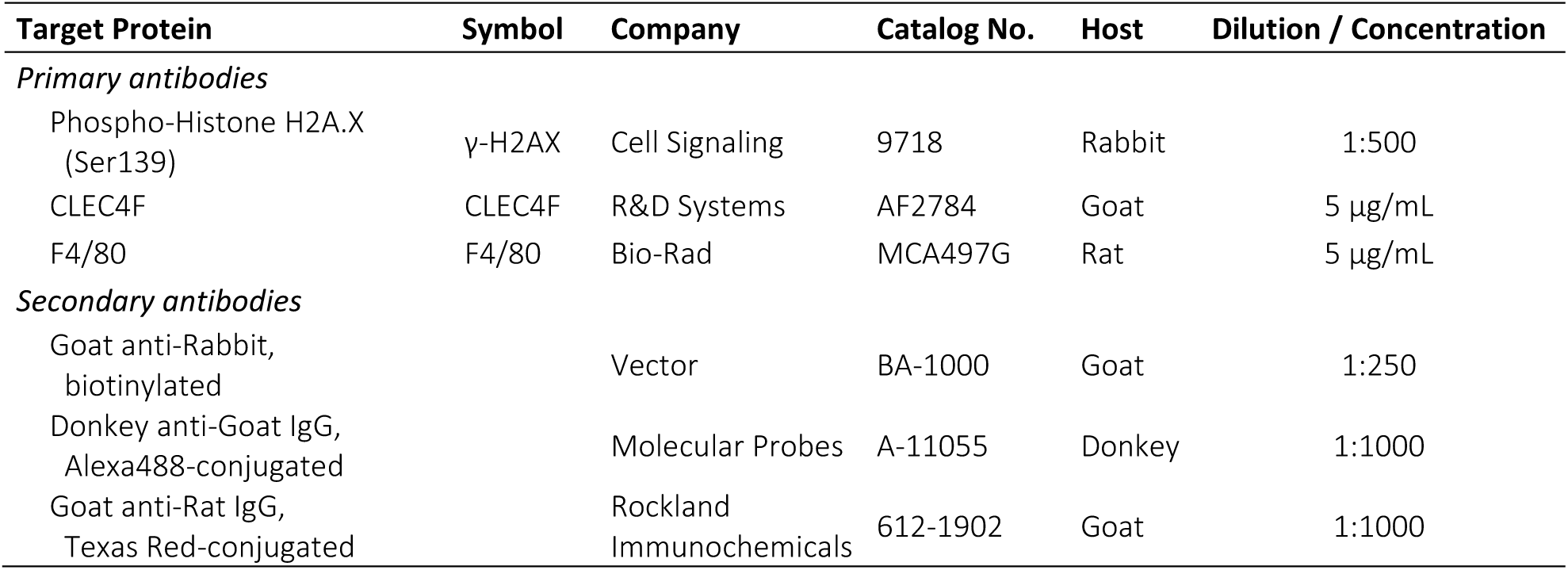
Primary antibodies for immunohistochemistry (IHC) and immunofluorescence (IF)

**Supplementary Table 2.** Primary antibodies for Western blot (WB)

| Target Protein | Symbol | Company | Catalog No. | Host | Dilution |
|---|---|---|---|---|---|
| *Primary antibodies* |  |  |  |  |  |
| Actin (all isoforms) | ACTN | abcam | ab3280 | Mouse | 1:1000 |
| cis-Aconitate decarboxylase | ACOD1/IRG1 | abcam | ab222411 | Rabbit | 1:1000 |
| Carnitine palmitoyltransferase I | CPT1A | abcam | ab234111 | Rabbit | 1:1000 |
| *Secondary antibodies* |  |  |  |  |  |
| Goat anti-mouse IgG, HRP-conjugated |  | Invitrogen | 31432 | Goat | 1:10000 |
| Goat anti-rabbit IgG, HRP-conjugated |  | Invitrogen | 31460 | Goat | 1:10000 |

**Supplementary Table 3.**
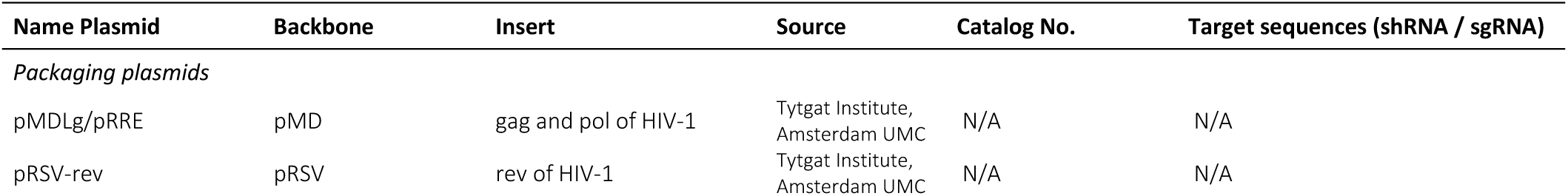
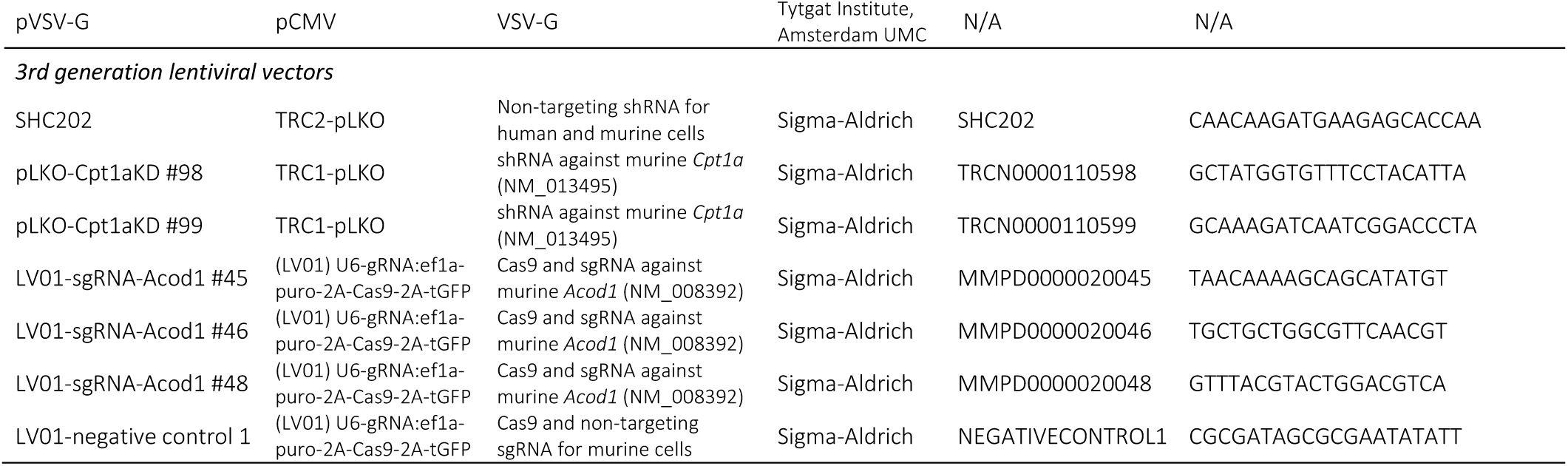
Lentivirus packaging plasmids and lentiviral vectors.

## 8. Supplementary Video Legends

**Video S1.** Three-dimensional maximum intensity projection (3D-MIP) rendering of HepOrgs co-cultured with HSCs under control conditions for 9 days. Lipidspot610 staining visualizes lipid droplets while Hoechst33342 staining highlights nuclei. (Related to Figure 1B)

**Video S2.** 3D-MIP rendering of HepOrgs co-cultured with HSCs under MASH conditions for 9 days. Lipidspot610 staining visualizes lipid droplets while Hoechst33342 staining highlights nuclei. (Related to Figure 1B)

Supplementary videos have been deposited in Zenodo (https://doi.org/10.5281/zenodo.24440966).

## 9. Supplementary Spreadsheets

**Spreadsheet S1.** Intracellular metabolites identified by LC-MS-based metabolomics with [U-^13^C6]-glucose tracing in the MASH-in-a-dish model. The spreadsheet contains both raw and natural abundance-corrected, normalized signal intensities (related to Figure 2 and Figure 3).

**Spreadsheet S2.** Extracellular (i.e. medium) metabolites identified by LC-MS-based metabolomics with [U-^13^C6]-glucose tracing in the MASH-in-a-dish model. The spreadsheet contains both raw and natural abundance-corrected signal intensities (related to Figure 2).

Supplementary spreadsheets have been deposited in Zenodo (https://doi.org/10.5281/zenodo.24440966).

